# Gramicidin S and Melittin - Potential anti-viral therapeutic peptides to treat SARS-CoV-2 infection

**DOI:** 10.1101/2021.10.21.465254

**Authors:** Mohammed Ghalib, Yash Parekh, Sarena Banu, Sushma Ram, Ramakrishnan Nagaraj, Bokara Kiran Kumar, Mohammed M Idris

## Abstract

The COVID19 pandemic has resulted in multipronged approaches for treatment of the disease. Since *de novo* discovery of drugs is time consuming, repurposing of molecules is now considered as one of the alternative strategies to treat COVID19. Antibacterial peptides are being recognized as attractive candidates for repurposing to treat viral infections. In this study, we describe the anti-SARS-CoV-2 activity of gramicidin S and melittin peptides obtained from *Bacillus brevis* and bee venom respectively. Our *in vitro* antiviral assay results showed significant decrease in the viral load compared to the untreated group with no/very less cytotoxicity. The EC_50_ values for gramicidin S and melittin are calculated as 1.571μg and 0.656μg respectively. Both the peptides treated to the SARS-CoV-2 infected Vero cells showed viral clearance from 12 hours onwards with a maximal clearance after 24 hours post infection. Based on proteome analysis it was found that more than 250 proteins were found to be differentially regulated in the gramicidin S and melittin treated SARS-CoV-2 infected Vero cells against control SARS-CoV-2 infected Vero cells after 24 and 48 hours post infection. The identified proteins were found to be associated in the metabolic and mRNA processing of the Vero cells post-treatment and infection. Both these peptides could be attractive candidates for repurposing to treat SARS-CoV-2 infection.

## Introduction

The pandemic caused by SARS-CoV-2 has led to intense research not only on the biology of the virus but also therapeutic intervention with a multi-pronged approach [1]. Vaccines have been developed at “warp” speed and have played a major role in controlling the disease [2]. However, vaccines are not available universally and there have been cases of infection with SARS-CoV-2 even in vaccinated individuals, though not severe [3]. There is clearly a need for development of therapeutic agents in addition to vaccines. In the area of anti-infective agents against SARS-CoV-2, efforts have been taken to generate therapeutic antibodies that would neutralize the virus and prevent its interaction with cellular receptors to gain entry into cells [4]. Considering the time scales in developing a drug *de novo*, there have been several attempts to re-purpose drugs to treat COVID19 [1, 5, 6]. However, repurposed drugs have had very limited success in treating SARS-CoV-2 infection including remdesivir [7]. There is no drug to-date that can be used specifically to treat COVID19 disease.

Infection in the case of SARS-Cov-2 is initiated by binding of the spike protein to ACE2 receptor followed by a series of steps leading to fusion and internalization of the virus and propagation [1, 8]. If the binding of the spike protein to ACE2 is prevented, then the virus will no longer be able to enter cells and propagate. SARS-CoV-2 is an enveloped virus where the RNA is encapsulated within a lipid vesicular structure with the spike protein decorating on the external side giving the “corona” appearance [1,8]. Disruption of the lipid structure would lead to the disintegration of the virus. Naturally occurring membrane-active peptides have potent antimicrobial activity which stems from their ability to disrupt bacterial membranes [9, 10]. We have explored the antiviral activity of two extensively studied peptides, gramicidin S having the sequence: [cyclo-(Val-Orn-Leu-D-Phe-Pro)2] [11] and the bee venom peptide melittin having the sequence: GIGAVLKVLTTGLPALISWIKRKRQQ-amide [12]. We reasoned that if the peptides could destabilize the viral membrane, the virus would disintegrate and would thus be rendered inactive. The peptides could also conceivably bind to the spike protein and prevents its interaction with ACE2 receptor or inhibit fusion. Peptides are crucial components of host-defense against bacteria and fungi in species across the evolutionary scale [9, 10]. There has been intense research in recent years to examine whether these and related peptides have the ability to neutralize SARS-CoV-2 and also have therapeutic potential [10, 13–15]. They include naturally occurring peptides that can be easily isolated. Many of these peptides have also been investigated structurally. These include gramicidin S and the bee venom peptide melittin [11, 12]. The antiviral activity of melittin against several viruses has been investigated extensively [16]. Nano-conjugates of melittin with sitagliptin have been investigated for anti-SARS-CoV-2 activity [17]. Other pharmacological activities of melittin have also been investigated [18, 19]. Gramicidin S has been used therapeutically to treat dental applications in humans [20]. We have investigated the antiviral activity of gramicidin S and melittin against SARS-CoV-2 in vitro in detail. We have observed that both the peptides have the ability to neutralize the virus in an *in vitro* assay using Vero cells. At the EC_50_ value, there is no cytolytic activity. The viral load had decreased drastically in the treatment group as seen by confocal microscopic images. Proteomic analysis indicates that there is also a metabolic effect and not merely viral lysis. Both the peptides could be attractive candidates for development as therapeutic agents to treat SARS-CoV-2 infection. As the viral membrane would be a likely target, mutant strains may likely be susceptible to the peptides.

## Results

### Antiviral activity of peptides

The SARS-CoV-2 viral particles enumerated by the RT-qPCR showed that treatment of melittin and gramicidin S effectively reduced viral load *in vitro* (Log EC_50_ value of gramicidin S (0.1963) corresponds to 1.571 μg and Log EC_50_ value (2.826) of melittin corresponds to 0.656 μg)Figure 1. The cytolytic activity of gramicidin S and melittin was determined using MTT assay (Figure 2). Results showed 75-80% cell survival at all the concentrations tested (up to 5 μg) indicating the safe use of these peptides. The antiviral activity of gramicidin S and melittin at 12 and 24 h was examined along with remdesivir as assay control. The data shown in Figure 3 indicates that the peptides show antiviral activity at 12 h and is more pronounced at 24 h. Log viral reduction is greater for the peptides at 24 h compared to remdesivir. The SARS-CoV-2 antiviral activity of gramicidin S and melittin was compared with remdesivir by confocal microscopy (Figure 4). Panel B (green fluorescence) indicates infection of cells, more prominent at 24 h. Panels D and E correspond to virus incubated with gramicidin S and melittin before incubating with cells. The considerable decrease in green fluorescence indicates that both the peptides have good anti-viral activity as with remdesivir shown in Panel E. Although, antiviral activity is observed at 12h, it is more pronounced at 24h.

**Figure 1:**
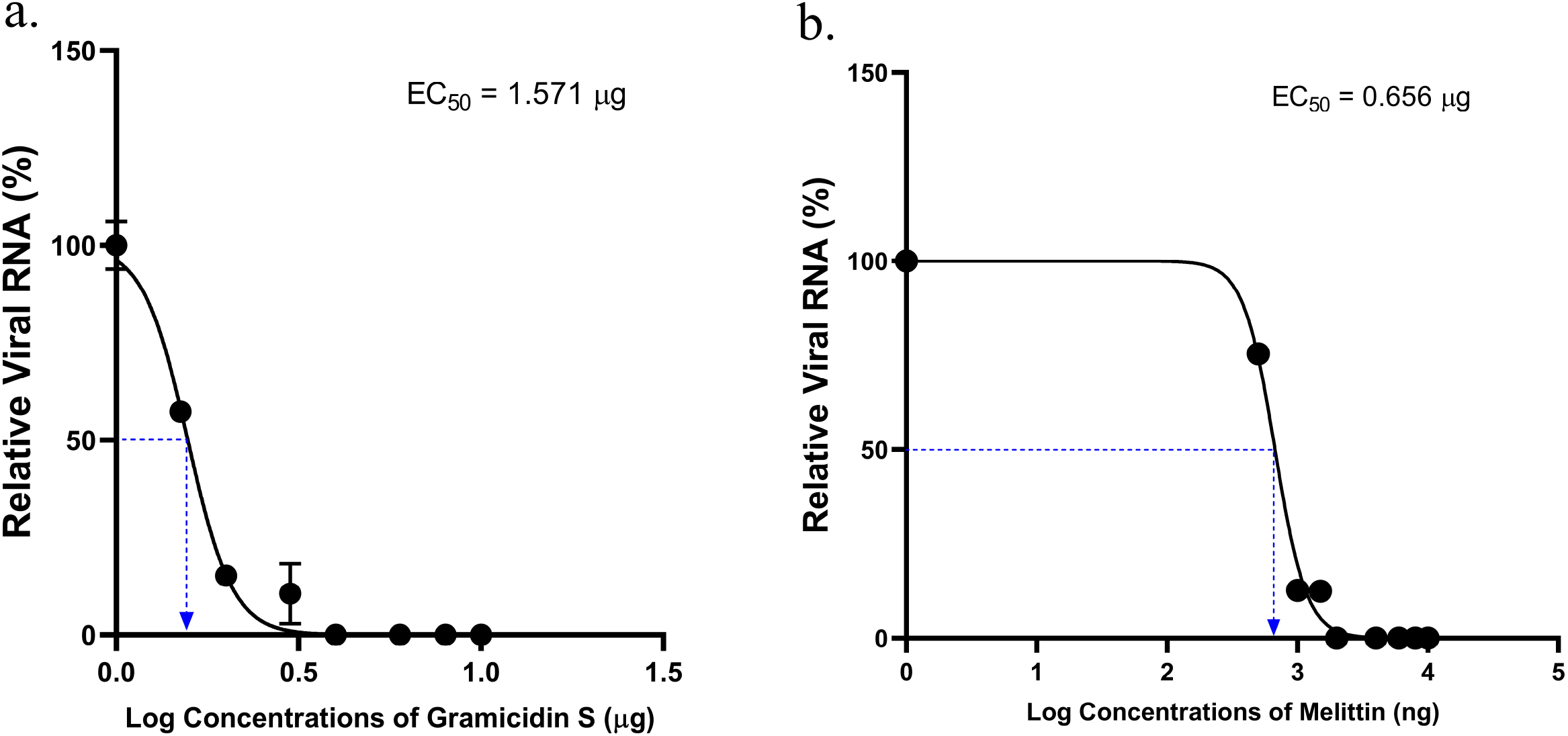
Anti SARS-CoV-2 activity of gramicidin S and melittin *in vitro*: a. Relative viral RNA (%) vs Log concentrations of Gramicidin S (μg). b.) Relative viral RNA (%) vs Log concentrations of melittin(ng). The graphs represent the Ct values of N-gene calculated using RT-qPCR in the supernatants. The Log EC_50_ value of gramicidin S (0.1963) corresponds to 1.571 μg and Log EC_50_ value (2.826) of melittincorresponds to 0.656 μg as shown in the graph

**Figure 2:**
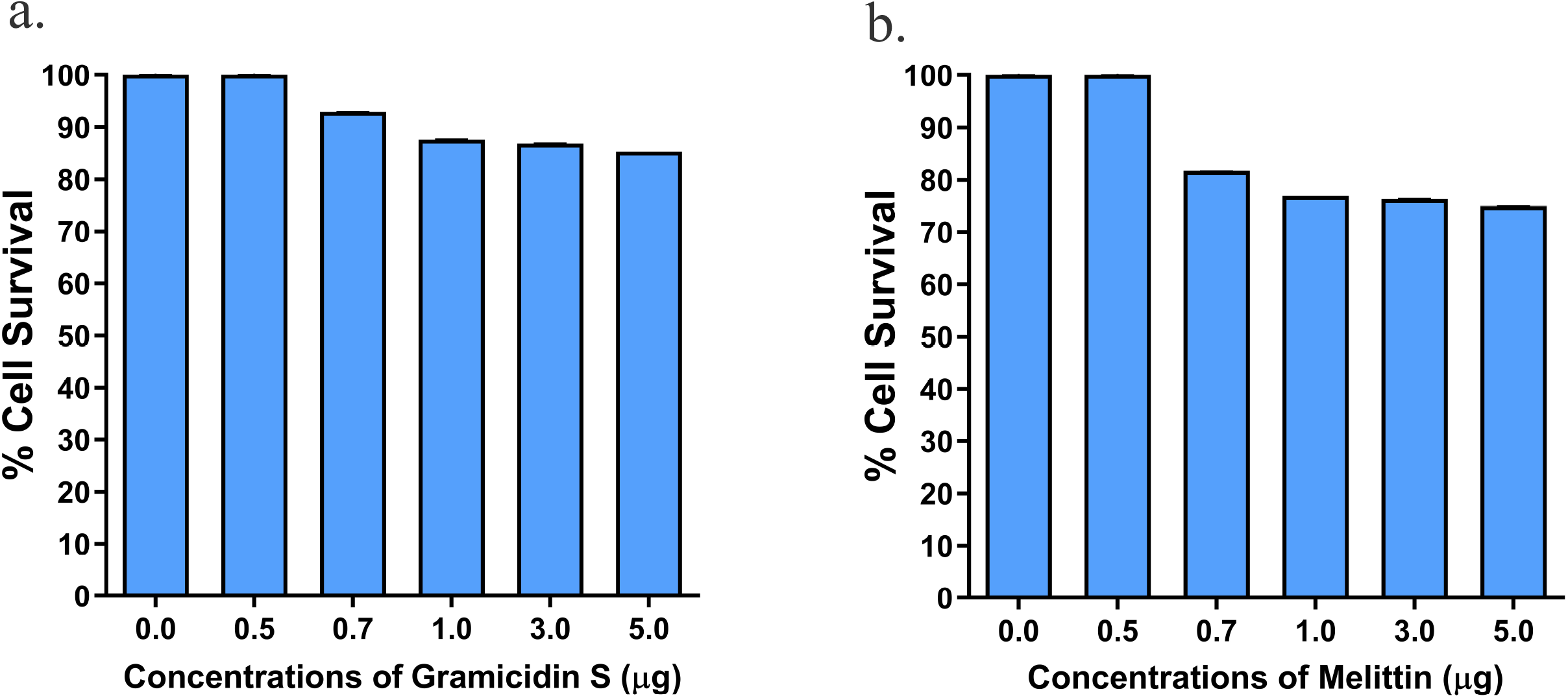
Measurement of Cytolytic activity of gramicidin S andmelittinusing MTT Assay: The graphs represent the percentage of cell viability vs concentrations of a. Gramicidin S (μg). bMelittin (μg).

**Figure 3:**
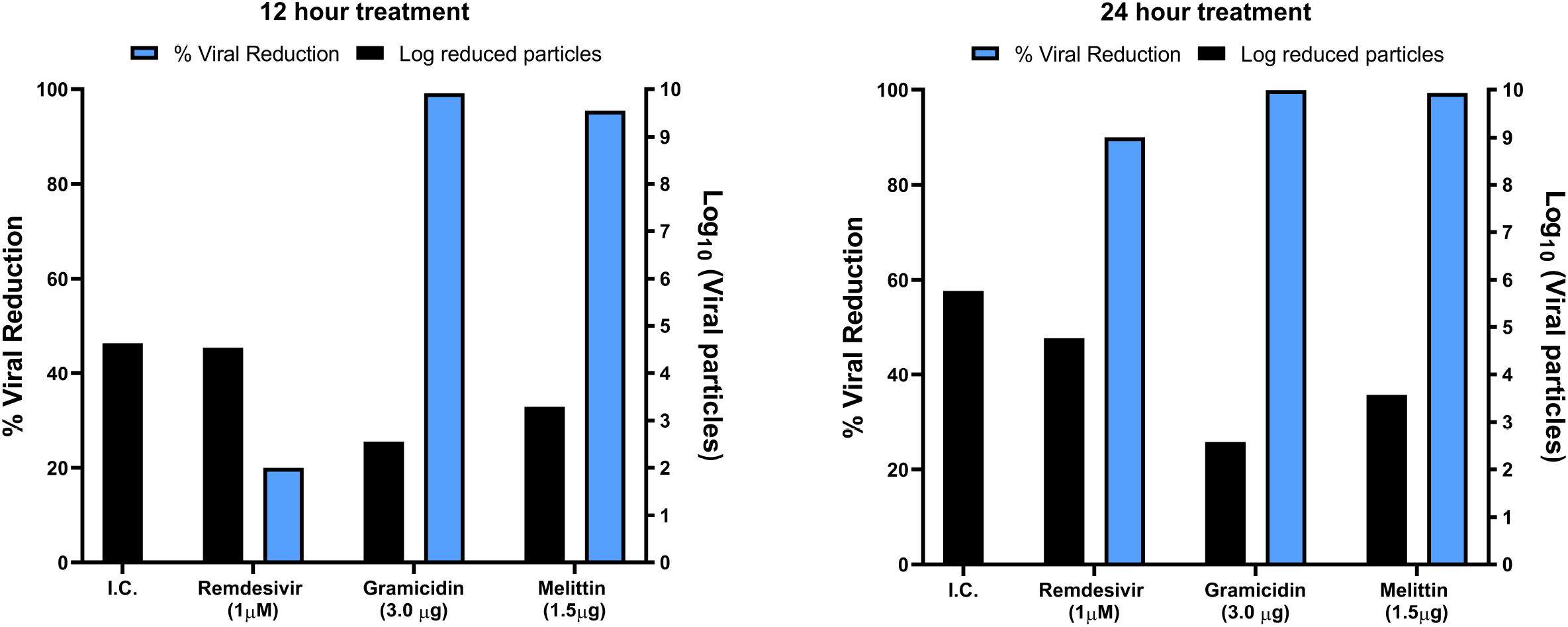
Gramicidin S (3.0 μg) and melittin (1.5 μg), were tested along with remdesivir(1.0 μM) (X-Axis) against SARS-CoV-2 *in vitro*. The graphs represent the % of viral reduction (Y-axis) and Log10 viral particles (Y’-Axis).

**Figure 4:**
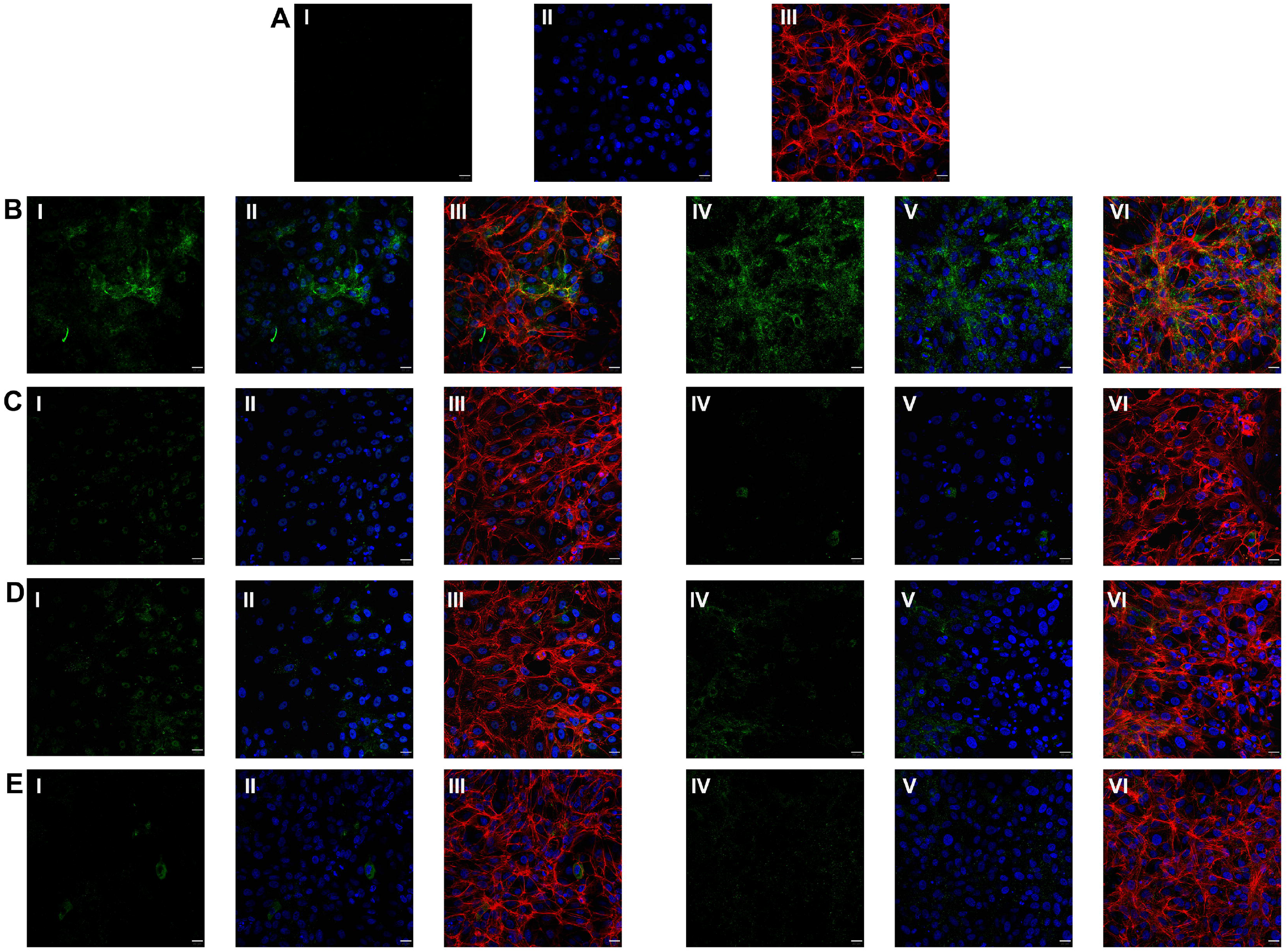
Immuno-fluorescence staining images against RBD protein expression specific to SARS-CoV-2 in Vero cells.(A) Vero cells without SARS-CoV-2 infection (mock)(I) no expression of RBD protein of SARS-CoV-2 virus; (II) DAPI staining representing nucleus (blue); (III) Merged image of f-Actin (Phalloidin staining, red), nucleus (DAPI, blue).(B) Vero cells infected with SARS-CoV-2.(I) represents RBD protein expression (green) of SARS-CoV-2 virus at 12 hours; (II) DAPI staining representing nucleus (blue) along with RBD protein expression of SARS-CoV-2 (green) at 12 hours; (III) Merged image of f-Actin (red) (Phalloidin staining), nucleus (DAPI) and RBD protein of SARS-CoV-2 (green) at 12 hours; (IV) represents RBD protein expression (green) of SARS-CoV-2 virus at 24 hours; (V) DAPI staining representing nucleus (blue) along with RBD protein expression of SARS-CoV-2 (green)at 24 hours; (VI) Merged image of f-Actin (Phalloidin staining, red), nucleus (DAPI, blue) and RBD protein of SARS-CoV-2 (green)at 24 hours.(C) Vero cells infected with SARS-CoV-2treated with gramicidin S.(I) represents RBD protein expression (green) of SARS-CoV-2 virus at 12 hours; (II) DAPI staining representing nucleus (blue) along with RBD protein expression of SARS-CoV-2 (green)at 12 hours; (III) Merged image of f-Actin (red) (Phalloidin staining), nucleus (DAPI) and RBD protein of SARS-CoV-2 (green)at 12 hours; (IV) represents RBD protein expression (green) of SARS-CoV-2 virus at 24 hours; (V) DAPI staining representing nucleus (blue) along with RBD protein expression of SARS-CoV-2 (green) at 24 hours; (VI) Merged image f-Actin (Phalloidin staining, red), nucleus (DAPI, blue) and RBD protein of SARS-CoV-2 (green) at 24 hours.((D)Vero cells infected with SARS-CoV-2 treated with melittin. (I) represents RBD protein expression (green) of SARS-CoV-2 virus at 12 hours; (II) DAPI staining representing nucleus (blue) along with RBD protein expression of SARS-CoV-2 (green) at 12 hours; (III) Merged image of f-Actin (red) (Phalloidin staining), nucleus (DAPI) and RBD protein of SARS-CoV-2 (green) at 12 hours; (IV) represents RBD protein expression (green) of SARS-CoV-2 virus at 24 hours; (V) DAPI staining representing nucleus (blue) along with RBD protein expression of SARS-CoV-2 (green) at 24 hours; (VI) Merged image of f-Actin (Phalloidin staining, red), nucleus (DAPI, blue) and RBD protein of SARS-CoV-2 (green) at 24 hours. E) Vero cells infected with SARS-CoV-2 treated with remdesivir.(I) represents RBD protein expression (green) of SARS-CoV-2 virus at 12 hours; (II) DAPI staining representing nucleus (blue) along with RBD protein expression of SARS-CoV-2 (green)at 12 hours; (III) Merged image of f-Actin (red) (Phalloidin staining), nucleus (DAPI) and RBD protein of SARS-CoV-2 (green) at 12 hours; (IV) represents RBD protein expression (green) of SARS-CoV-2 virus at 24 hours; (V) DAPI staining representing nucleus (blue) along with RBD protein expression of SARS-CoV-2 (green)at 24 hours; (VI) Merged image of f-Actin (Phalloidin staining, red), nucleus (DAPI, blue) and RBD protein of SARS-CoV-2 (green)at 24 hours. Scale bars, 20 μm (40X image).

### Proteomics Analysis

The effect of the peptides on the ability of virus to infect Vero cells was studied by proteomics. iTRAQ based quantitative proteomics analysis identified 7 SARS-CoV-2 proteins as up regulated in the control Vero Cells. Nsp9, ORF1ab, ORF10 and nucleocapsid phosphoprotein were found to be up-regulated in the Vero cells after 24 hours of infection, whereas the same proteins were found to be down regulated in the cells upon gramicidin S and melittin treatment (Table 1). Similarly, at 48 hours post infection (hpi), ORF1ab, S protein, nucleocapsid phosphoprotein, helicase and RNA-dependent RNA polymerase were found to be up-regulated in the infected Vero cells which were found to be down regulated in the gramicidin S and melittin treated Vero cells (Table 1).

**Table 1:** List of SARS-CoV-2 proteins and their expression level in the vero cells post infection (control), Melittin treatment and Gramicidin treatment.

| Time Point | Protein Accession No. | Protein Name | # Peptides | # PSMs | Infected Controls | Gramicidin S treated Infected cells | Melittin treated Infected cells |
| --- | --- | --- | --- | --- | --- | --- | --- |
| 24hpi | YP_009742616.1 | nsp9 [Severe acute respiratory syndrome coronavirus 2] | 17 | 69 | 0.1 | -1.2 | -0.9 |
|  | YP_009724389.1 | ORF1ab polyprotein [Severe acute respiratory syndrome coronavirus 2] | 762 | 3246 | 0.3 | -1.3 | -1.1 |
|  | YP_009725255.1 | ORF10 protein [Severe acute respiratory syndrome coronavirus 2] | 4 | 9 | 0.5 | -1.2 | -1.4 |
|  | YP_009724397.2 | nucleocapsid phosphoprotein [Severe acute respiratory syndrome coronavirus 2] | 71 | 260 | 0.6 | 0.1 | 1.6 |
| 48hpi | YP_009724389.1 | ORF1ab polyprotein [Severe acute respiratory syndrome coronavirus 2] | 755 | 3587 | 0.8 | -0.8 | -0.9 |
|  | YP_009724390.1 | surface glycoprotein [Severe acute respiratory syndrome coronavirus 2] | 108 | 418 | 0.1 | -1.3 | -0.7 |
|  | YP_009724397.2 | nucleocapsid phosphoprotein [Severe acute respiratory syndrome coronavirus 2] | 75 | 271 | 2.2 | -2.8 | -2.7 |
|  | YP_009725308.1 | helicase [Severe acute respiratory syndrome coronavirus 2] | 83 | 312 | 0.2 | -1.2 | -0.3 |
|  | YP_009725307.1 | RNA-dependent RNA polymerase [Severe acute respiratory syndrome coronavirus 2] | 106 | 492 | 0.9 | -1.0 | -1.6 |

Based on proteomic analysis it was found that a total of 254 proteins were found to be associated with differentially regulated in Vero cells (Table 2 and Figure 5a). It was found that majority of down and up-regulated proteins were upturned with gramicidin S and melittin treatment at 24 and 48 hpi. RS28, K22E, K2C1, RL17 are few of the major down-regulated proteins which were found to be reversing their expression post peptide treatment. NPM, ACLY, CALX and F184B were found to be up-regulated in Vero cells 24hpi, whereas peptide treatment showed reversal of the protein expression (Table 2). Based on heat map and cluster analysis it is very interesting to see the tight association of peptide treated Vero cell protein expression against control infected Vero cell protein expression for both the time points post infection (Figure5a).

**Table 2:** List of proteins expressed in the vero cells post infection, infection and melittin treatment and infection and Gramicidin S treatment for 24 and 48 hours post infection.

| UNIPROT ID | Description | Symbol | # Peptides | # PSMs | 24 hours post infection (hpi) |  |  | 48 hours post infection (hpi) |  |  |
| --- | --- | --- | --- | --- | --- | --- | --- | --- | --- | --- |
|  |  |  |  |  | Infected controls | Gramicidin S treated Infected cells | Melittin treated Infected cells | Infected controls | Gramicidin S treated Infected cells | Melittin treated Infected cells |
| O14744 | Protein arginine N-methyltransferase 5 OS=Homo sapiens OX=9606 GN=PRMT5 PE=1 SV=4 - [ANM5_HUMAN] | ANM5 | 7 | 9 | 0.1 | -1.4 | -0.9 | -0.9 | -0.9 | 0.7 |
| O14818 | Proteasome subunit alpha type-7 OS=Homo sapiens OX=9606 GN=PSMA7 PE=1 SV=1 - [PSA7_HUMAN] | PSA7 | 5 | 7 | -0.3 | -1.8 | -0.7 | 0.2 | -0.1 | 0.1 |
| O14979 | Heterogeneous nuclear ribonucleoprotein D-like OS=Homo sapiens OX=9606 GN=HNRNPDL PE=1 SV=3 - [HNRDL_HUMAN] | HNRDL | 6 | 9 | 0.4 | -1.8 | -1.0 | 0.1 | -1.5 | 0.2 |
| O43175 | D-3-phosphoglycerate dehydrogenase OS=Homo sapiens OX=9606 GN=PHGDH PE=1 SV=4 - [SERA_HUMAN] | SERA | 5 | 6 | -0.4 | -1.0 | -0.3 | 1.0 | -0.3 | -1.2 |
| O60218 | Aldo-keto reductase family 1 member B10 OS=Homo sapiens OX=9606 GN=AKR1B10 PE=1 SV=2 - [AK1BA_HUMAN] | AK1BA | 6 | 7 | 1.4 | -0.8 | 1.5 | -0.9 | -0.2 | 0.2 |
| O60318 | Germinal-center associated nuclear protein OS=Homo sapiens OX=9606 GN=MCM3AP PE=1 SV=2 - [GANP_HUMAN] | GANP | 16 | 18 | -0.4 | -1.0 | -0.3 | 0.1 | -1.6 | 0.0 |
| O60701 | UDP-glucose 6-dehydrogenase OS=Homo sapiens OX=9606 GN=UGDH PE=1 SV=1 - [UGDH_HUMAN] | UGDH | 8 | 8 | -0.5 | -1.3 | -0.7 | 1.6 | -0.4 | -2.3 |
| O75390 | Citrate synthase, mitochondrial OS=Homo sapiens OX=9606 GN=CS PE=1 SV=2 - [CISY_HUMAN] | CISY | 8 | 10 | 1.4 | -0.6 | 1.4 | -1.1 | -1.0 | -0.7 |
| O95359 | Transforming acidic coiled-coil-containing protein 2 OS=Homo sapiens OX=9606 GN=TACC2 PE=1 SV=3 - [TACC2_HUMAN] | TACC2 | 14 | 18 | -1.0 | -1.4 | -1.7 | 1.3 | -1.0 | -1.5 |
| O95831 | Apoptosis-inducing factor 1, mitochondrial OS=Homo sapiens OX=9606 GN=AIFM1 PE=1 SV=1 - [AIFM1_HUMAN] | AIFM1 | 9 | 11 | 0.1 | -2.0 | -0.9 | 1.0 | -0.6 | -1.5 |
| P00338 | L-lactate dehydrogenase A chain OS=Homo sapiens OX=9606 GN=LDHA PE=1 SV=2 - [LDHA_HUMAN] | LDHA | 14 | 29 | 1.3 | -1.2 | 0.7 | -1.5 | -1.6 | -1.7 |
| P00352 | Retinal dehydrogenase 1 OS=Homo sapiens OX=9606 GN=ALDH1A1 PE=1 SV=2 - [AL1A1_HUMAN] | AL1A1 | 5 | 9 | 0.3 | -1.5 | -1.5 | -0.4 | -1.2 | 0.0 |
| P00390 | Glutathione reductase, mitochondrial OS=Homo sapiens OX=9606 GN=GSR PE=1 SV=2 - [GSHR_HUMAN] | GSHR | 7 | 10 | 0.2 | -1.9 | -1.5 | -0.6 | -0.9 | 0.3 |
| P00492 | Hypoxanthine-guanine phosphoribosyltransferase OS=Homo sapiens OX=9606 GN=HPRT1 PE=1 SV=2 - [HPRT_HUMAN] | HPRT | 1 | 1 | 0.0 | -1.3 | -1.0 | 0.4 | -2.3 | -0.9 |
| P00505 | Aspartate aminotransferase, mitochondrial OS=Homo sapiens OX=9606 GN=GOT2 PE=1 SV=3 - [AATM_HUMAN] | AATM | 8 | 11 | 1.1 | -0.7 | 1.0 | -1.9 | -1.8 | -1.5 |
| P00558 | Phosphoglycerate kinase 1 OS=Homo sapiens OX=9606 GN=PGK1 PE=1 SV=3 - [PGK1_HUMAN] | PGK1 | 19 | 33 | 1.4 | -0.7 | 1.6 | -1.6 | -1.8 | -2.2 |
| P00966 | Argininosuccinate synthase OS=Homo sapiens OX=9606 GN=ASS1 PE=1 SV=2 - [ASSY_HUMAN] | ASSY | 6 | 6 | 0.7 | -1.3 | 0.6 | -1.3 | -0.3 | 0.6 |
| P02545 | Prelamin-A/C OS=Homo sapiens OX=9606 GN=LMNA PE=1 SV=1 - [LMNA_HUMAN] | LMNA | 17 | 17 | -0.9 | -1.4 | -0.9 | 1.6 | -1.7 | -0.5 |
| P02751 | Fibronectin OS=Homo sapiens OX=9606 GN=FN1 PE=1 SV=4 - [FINC_HUMAN] | FINC | 18 | 24 | -1.2 | -1.2 | 0.6 | -0.2 | -0.8 | -1.3 |
| P02768 | Serum albumin OS=Homo sapiens OX=9606 GN=ALB PE=1 SV=2 - [ALBU_HUMAN] | ALBU | 9 | 15 | -1.7 | 1.1 | -0.3 | 0.5 | 0.5 | -1.7 |
| P04075 | Fructose-bisphosphate aldolase A OS=Homo sapiens OX=9606 GN=ALDOA PE=1 SV=2 - [ALDOA_HUMAN] | ALDOA | 21 | 36 | 1.2 | -0.6 | 1.3 | -1.7 | -1.6 | -1.3 |
| P04083 | Annexin A1 OS=Homo sapiens OX=9606 GN=ANXA1 PE=1 SV=2 - [ANXA1_HUMAN] | ANXA1 | 18 | 36 | 0.6 | -0.4 | 1.4 | -1.4 | -1.2 | -1.7 |
| P04179 | Superoxide dismutase [Mn], mitochondrial OS=Homo sapiens OX=9606 GN=SOD2 PE=1 SV=3 - [SODM_HUMAN] | SOD2 | 6 | 12 | 0.4 | 0.6 | 0.7 | 2.2 | -1.1 | -1.5 |
| P04264 | Keratin, type II cytoskeletal 1 OS=Homo sapiens OX=9606 GN=KRT1 PE=1 SV=6 - [K2C1_HUMAN] | K2C1 | 32 | 63 | -2.6 | 0.1 | 0.5 | -0.6 | -0.9 | -0.7 |
| P04406 | Glyceraldehyde-3-phosphate dehydrogenase OS=Homo sapiens OX=9606 GN=GAPDH PE=1 SV=3 - [G3P_HUMAN] | G3P | 19 | 67 | 1.4 | -0.9 | 1.4 | -1.9 | -1.5 | -1.9 |
| P04792 | Heat shock protein beta-1 OS=Homo sapiens OX=9606 GN=HSPB1 PE=1 SV=2 - [HSPB1_HUMAN] | HSPB1 | 11 | 21 | -1.1 | -1.9 | -0.6 | 0.0 | -0.2 | -0.1 |
| P05121 | Plasminogen activator inhibitor 1 OS=Homo sapiens OX=9606 GN=SERPINE1 PE=1 SV=1 - [PAI1_HUMAN] | PAI1 | 5 | 6 | -0.5 | -0.6 | -0.1 | -0.5 | -1.1 | 0.1 |
| P05141 | ADP/ATP translocase 2 OS=Homo sapiens OX=9606 GN=SLC25A5 PE=1 SV=7 - [ADT2_HUMAN] | ADT2 | 13 | 22 | -0.7 | -0.5 | -0.6 | -0.6 | -0.8 | 0.2 |
| P05386 | 60S acidic ribosomal protein P1 OS=Homo sapiens OX=9606 GN=RPLP1 PE=1 SV=1 - [RLA1_HUMAN] | RLA1 | 1 | 7 | -1.9 | -1.2 | 0.9 | 2.6 | -1.8 | -2.7 |
| P05388 | 60S acidic ribosomal protein P0 OS=Homo sapiens OX=9606 GN=RPLP0 PE=1 SV=1 - [RLA0_HUMAN] | RLA0 | 10 | 24 | 1.2 | -0.2 | 1.7 | -1.4 | -1.6 | -2.0 |
| P05783 | Keratin, type I cytoskeletal 18 OS=Homo sapiens OX=9606 GN=KRT18 PE=1 SV=2 - [K1C18_HUMAN] | K1C18 | 15 | 20 | -0.7 | -1.3 | -0.8 | -2.1 | -1.2 | -1.2 |
| P06576 | ATP synthase subunit beta, mitochondrial OS=Homo sapiens OX=9606 GN=ATP5F1B PE=1 SV=3 - [ATPB_HUMAN] | ATPB | 11 | 20 | 0.3 | -1.8 | -1.1 | 1.3 | -0.7 | -1.6 |
| P06733 | Alpha-enolase OS=Homo sapiens OX=9606 GN=ENO1 PE=1 SV=2 - [ENOA_HUMAN] | ENOA | 23 | 45 | -0.6 | -1.6 | -0.4 | 1.2 | -0.5 | -1.9 |
| P06744 | Glucose-6-phosphate isomerase OS=Homo sapiens OX=9606 GN=GPI PE=1 SV=4 - [G6PI_HUMAN] | G6PI | 6 | 10 | -0.1 | 0.0 | 1.9 | 2.0 | -1.8 | -0.8 |
| P06748 | Nucleophosmin OS=Homo sapiens OX=9606 GN=NPM1 PE=1 SV=2 - [NPM_HUMAN] | NPM | 18 | 38 | 2.2 | -1.1 | 1.5 | -2.1 | -0.9 | -0.8 |
| P06753 | Tropomyosin alpha-3 chain OS=Homo sapiens OX=9606 GN=TPM3 PE=1 SV=2 - [TPM3_HUMAN] | TPM3 | 13 | 22 | 1.3 | -1.2 | 0.8 | -1.5 | -1.0 | -1.1 |
| P07195 | L-lactate dehydrogenase B chain OS=Homo sapiens<br>OX=9606 GN=LDHB PE=1 SV=2 - [LDHB_HUMAN] | LDHB | 16 | 22 | 1.5 | -1.1 | 1.1 | -1.9 | -1.2 | -1.4 |
| P07237 | Protein disulfide-isomerase OS=Homo sapiens<br>OX=9606 GN=P4HB PE=1 SV=3 - [PDIA1_HUMAN] | PDIA1 | 13 | 17 | 0.0 | 0.1 | 2.7 | -0.2 | -1.5 | 0.1 |
| P07305 | Histone H1.0 OS=Homo sapiens OX=9606 GN=H1F0<br>PE=1 SV=3 - [H10_HUMAN] | H10 | 1 | 2 | 0.2 | -0.4 | -0.4 | 0.2 | -1.8 | -0.8 |
| P07355 | Annexin A2 OS=Homo sapiens OX=9606 GN=ANXA2<br>PE=1 SV=2 - [ANXA2_HUMAN] | ANXA2 | 37 | 100 | 1.4 | -0.6 | 1.4 | -2.0 | -1.2 | -1.6 |
| P07437 | Tubulin beta chain OS=Homo sapiens OX=9606<br>GN=TUBB PE=1 SV=2 - [TBB5_HUMAN] | TBB5 | 21 | 48 | -0.5 | -1.8 | -0.8 | 1.3 | -0.6 | -2.2 |
| P07711 | Cathepsin L1 OS=Homo sapiens OX=9606 GN=CTSL<br>PE=1 SV=2 - [CATL1_HUMAN] | CATL1 | 8 | 11 | -0.2 | -1.6 | -0.8 | -0.1 | 0.3 | 0.2 |
| P07737 | Profilin-1 OS=Homo sapiens OX=9606 GN=PFN1<br>PE=1 SV=2 - [PROF1_HUMAN] | PROF1 | 4 | 6 | 0.9 | 0.7 | 0.9 | 2.3 | -1.9 | -3.1 |
| P07910 | Heterogeneous nuclear ribonucleoproteins C1/C2<br>OS=Homo sapiens OX=9606 GN=HNRNPC PE=1<br>SV=4 - [HNRPC_HUMAN] | HNRPC | 12 | 18 | 1.5 | -1.1 | 0.9 | -2.1 | -1.5 | -1.6 |
| P08133 | Annexin A6 OS=Homo sapiens OX=9606 GN=ANXA6<br>PE=1 SV=3 - [ANXA6_HUMAN] | ANXA6 | 12 | 14 | -0.6 | -0.8 | -1.1 | -0.3 | -1.1 | 0.1 |
| P08238 | Heat shock protein HSP 90-beta OS=Homo sapiens<br>OX=9606 GN=HSP90AB1 PE=1 SV=4 -<br>[HS90B_HUMAN] | HS90B | 30 | 60 | 0.3 | -1.4 | -1.1 | -0.2 | 0.5 | -1.1 |
| P08243 | Asparagine synthetase [glutamine-hydrolyzing]<br>OS=Homo sapiens OX=9606 GN=ASNS PE=1 SV=4 -<br>[ASNS_HUMAN] | ASNS | 4 | 4 | 0.0 | -1.9 | -0.9 | -1.4 | -0.3 | 0.9 |
| P08670 | Vimentin OS=Homo sapiens OX=9606 GN=VIM PE=1<br>SV=4 - [VIME_HUMAN] | VIME | 35 | 65 | -0.9 | -1.2 | -0.7 | 1.3 | -0.5 | -1.6 |
| P08758 | Annexin A5 OS=Homo sapiens OX=9606 GN=ANXA5<br>PE=1 SV=2 - [ANXA5_HUMAN] | ANXA5 | 14 | 24 | -0.6 | -0.9 | -0.2 | -1.2 | -1.2 | -1.4 |
| P08865 | 40S ribosomal protein SA OS=Homo sapiens OX=9606<br>GN=RPSA PE=1 SV=4 - [RSSA_HUMAN] | RSSA | 7 | 14 | 0.4 | 0.2 | 1.5 | -1.2 | -0.8 | -2.8 |
| P09211 | Glutathione S-transferase P OS=Homo sapiens<br>OX=9606 GN=GSTP1 PE=1 SV=2 -<br>[GSTP1_HUMAN] | GSTP1 | 2 | 2 | -2.0 | -2.0 | 0.9 | 4.7 | -2.1 | -3.5 |
| P09382 | Galectin-1 OS=Homo sapiens OX=9606 GN=LGALS1<br>PE=1 SV=2 - [LEG1_HUMAN] | LEG1 | 4 | 7 | -1.8 | -0.9 | 0.7 | 1.8 | -1.9 | -2.4 |
| P09525 | Annexin A4 OS=Homo sapiens OX=9606 GN=ANXA4<br>PE=1 SV=4 - [ANXA4_HUMAN] | ANXA4 | 15 | 26 | 0.5 | -0.6 | 1.0 | 0.5 | -1.0 | -0.4 |
| P09651 | Heterogeneous nuclear ribonucleoprotein A1 OS=Homo<br>sapiens OX=9606 GN=HNRNPA1 PE=1 SV=5 -<br>[ROA1_HUMAN] | ROA1 | 21 | 45 | -0.7 | -2.5 | -0.8 | -0.1 | -0.3 | 0.2 |
| P09972 | Fructose-bisphosphate aldolase C OS=Homo sapiens<br>OX=9606 GN=ALDOC PE=1 SV=2 -<br>[ALDOC_HUMAN] | ALDOC | 6 | 15 | -0.4 | -1.1 | -1.5 | -2.1 | -1.0 | -1.2 |
| P0CG48 | Polyubiquitin-C OS=Homo sapiens OX=9606 GN=UBC<br>PE=1 SV=3 - [UBC_HUMAN] | UBC | 8 | 10 | 1.1 | 0.2 | 1.3 | 2.6 | -1.6 | -1.9 |
| P10412 | Histone H1.4 OS=Homo sapiens OX=9606<br>GN=HIST1H1E PE=1 SV=2 - [H14_HUMAN] | H14 | 13 | 18 | 1.1 | -0.1 | 1.9 | 0.5 | -1.9 | -2.3 |
| P10809 | 60 kDa heat shock protein, mitochondrial OS=Homo sapiens OX=9606 GN=HSPD1 PE=1 SV=2 - [CH60_HUMAN] | CH60 | 20 | 24 | -1.0 | 1.1 | 3.2 | 1.2 | -0.5 | -2.0 |
| P11021 | Endoplasmic reticulum chaperone BiP OS=Homo sapiens OX=9606 GN=HSPA5 PE=1 SV=2 - [BIP_HUMAN] | BIP | 17 | 35 | -0.8 | -0.9 | 0.1 | 0.4 | -0.2 | -0.9 |
| P11142 | Heat shock cognate 71 kDa protein OS=Homo sapiens OX=9606 GN=HSPA8 PE=1 SV=1 - [HSP7C_HUMAN] | HSP7C | 22 | 31 | -0.1 | 0.2 | 2.4 | 1.6 | -1.2 | -1.8 |
| P11413 | Glucose-6-phosphate 1-dehydrogenase OS=Homo sapiens OX=9606 GN=G6PD PE=1 SV=4 - [G6PD_HUMAN] | G6PD | 6 | 12 | 0.0 | -2.1 | -0.9 | -2.0 | 0.4 | 1.2 |
| P11586 | C-1-tetrahydrofolate synthase, cytoplasmic OS=Homo sapiens OX=9606 GN=MTHFD1 PE=1 SV=3 - [C1TC_HUMAN] | C1TC | 10 | 11 | 0.4 | -0.8 | -0.9 | 0.1 | 0.4 | -0.4 |
| P12236 | ADP/ATP translocase 3 OS=Homo sapiens OX=9606 GN=SLC25A6 PE=1 SV=4 - [ADT3_HUMAN] | ADT3 | 5 | 9 | -0.2 | -0.8 | -1.4 | -0.1 | 0.5 | 0.2 |
| P13489 | Ribonuclease inhibitor OS=Homo sapiens OX=9606 GN=RNH1 PE=1 SV=2 - [RINI_HUMAN] | RINI | 3 | 6 | 1.6 | -1.2 | 0.8 | -1.7 | 0.0 | 0.8 |
| P13639 | Elongation factor 2 OS=Homo sapiens OX=9606 GN=EEF2 PE=1 SV=4 - [EF2_HUMAN] | EF2 | 23 | 35 | 0.3 | -1.3 | -1.0 | -2.3 | 1.3 | 2.4 |
| P13645 | Keratin, type I cytoskeletal 10 OS=Homo sapiens OX=9606 GN=KRT10 PE=1 SV=6 - [K1C10_HUMAN] | K1C10 | 22 | 41 | -0.9 | -1.0 | -1.0 | -0.3 | -1.4 | -1.1 |
| P13647 | Keratin, type II cytoskeletal 5 OS=Homo sapiens OX=9606 GN=KRT5 PE=1 SV=3 - [K2C5_HUMAN] | K2C5 | 15 | 17 | -0.9 | 0.1 | 2.4 | 1.8 | -0.4 | -2.4 |
| P13667 | Protein disulfide-isomerase A4 OS=Homo sapiens OX=9606 GN=PDIA4 PE=1 SV=2 - [PDIA4_HUMAN] | PDIA4 | 7 | 8 | 0.6 | -1.9 | -1.4 | -0.4 | -4.0 | -1.1 |
| P14618 | Pyruvate kinase PKM OS=Homo sapiens OX=9606 GN=PKM PE=1 SV=4 - [KPYM_HUMAN] | KPYM | 28 | 44 | -0.7 | 0.2 | 2.3 | 1.6 | -0.8 | -2.5 |
| P14625 | Endoplasmic reticulum chaperone OS=Homo sapiens OX=9606 GN=HSP90B1 PE=1 SV=1 - [ENPL_HUMAN] | ENPL | 20 | 31 | 0.4 | -1.4 | -1.3 | -3.3 | -0.9 | -1.1 |
| P14868 | Aspartate--tRNA ligase, cytoplasmic OS=Homo sapiens OX=9606 GN=DARS PE=1 SV=2 - [SYDC_HUMAN] | SYDC | 8 | 9 | 0.0 | -2.1 | -0.7 | -1.7 | 0.1 | 1.0 |
| P15121 | Aldose reductase OS=Homo sapiens OX=9606 GN=AKR1B1 PE=1 SV=3 - [ALDR_HUMAN] | ALDR | 14 | 29 | 1.3 | -1.5 | 0.6 | -2.1 | -1.1 | -1.2 |
| P15259 | Phosphoglycerate mutase 2 OS=Homo sapiens OX=9606 GN=PGAM2 PE=1 SV=3 - [PGAM2_HUMAN] | PGAM2 | 6 | 7 | 0.3 | 0.6 | 0.9 | 0.0 | -1.6 | -0.7 |
| P15559 | NAD(P)H dehydrogenase [quinone] 1 OS=Homo sapiens OX=9606 GN=NQO1 PE=1 SV=1 - [NQO1_HUMAN] | NQO1 | 3 | 4 | -0.2 | -1.4 | -1.4 | -0.1 | 0.1 | -0.2 |
| P15880 | 40S ribosomal protein S2 OS=Homo sapiens OX=9606 GN=RPS2 PE=1 SV=2 - [RS2_HUMAN] | RS2 | 14 | 17 | -0.6 | -1.3 | -0.3 | -0.4 | 0.4 | 0.3 |
| P16152 | Carbonyl reductase [NADPH] 1 OS=Homo sapiens OX=9606 GN=CBR1 PE=1 SV=3 - [CBR1_HUMAN] | NADPH | 8 | 11 | 1.2 | -0.4 | 1.7 | 0.0 | 0.2 | 0.1 |
| P17844 | Probable ATP-dependent RNA helicase DDX5 OS=Homo sapiens OX=9606 GN=DDX5 PE=1 SV=1 - [DDX5_HUMAN] | DDX5 | 12 | 15 | -0.9 | 0.3 | 2.2 | 0.4 | -1.9 | -0.1 |
| P17987 | T-complex protein 1 subunit alpha OS=Homo sapiens<br>OX=9606 GN=TCP1 PE=1 SV=1 - [TCPA_HUMAN] | TCPA | 6 | 7 | 0.1 | -1.8 | -1.1 | 1.4 | -2.9 | -1.2 |
| P18124 | 60S ribosomal protein L7 OS=Homo sapiens OX=9606<br>GN=RPL7 PE=1 SV=1 - [RL7_HUMAN] | RL7 | 9 | 11 | 0.0 | 0.5 | 1.3 | 1.1 | -1.8 | -1.4 |
| P18621 | 60S ribosomal protein L17 OS=Homo sapiens OX=9606<br>GN=RPL17 PE=1 SV=3 - [RL17_HUMAN] | RL17 | 3 | 4 | -2.0 | -1.5 | 0.3 | -2.7 | 1.7 | 4.2 |
| P18669 | Phosphoglycerate mutase 1 OS=Homo sapiens<br>OX=9606 GN=PGAM1 PE=1 SV=2 - [PGAM1_HUMAN] | PGAM1 | 6 | 7 | -0.4 | -1.5 | -1.2 | -0.8 | 0.4 | 0.7 |
| P19105 | Myosin regulatory light chain 12A OS=Homo sapiens<br>OX=9606 GN=MYL12A PE=1 SV=2 - [ML12A_HUMAN] | ML12A | 4 | 8 | -0.6 | -1.2 | -0.2 | 1.0 | -1.0 | -1.8 |
| P19338 | Nucleolin OS=Homo sapiens OX=9606 GN=NCL PE=1<br>SV=3 - [NUCL_HUMAN] | NUCL | 11 | 14 | 0.5 | -1.3 | -1.1 | 0.2 | 0.6 | -0.8 |
| P20073 | Annexin A7 OS=Homo sapiens OX=9606 GN=ANXA7<br>PE=1 SV=3 - [ANXA7_HUMAN] | ANXA7 | 6 | 7 | 0.1 | -1.6 | -1.2 | -1.0 | -2.1 | -1.7 |
| P21291 | Cysteine and glycine-rich protein 1 OS=Homo sapiens<br>OX=9606 GN=CSRP1 PE=1 SV=3 - [CSRP1_HUMAN] | CSRP1 | 5 | 7 | -0.8 | -0.9 | -0.8 | -1.6 | 0.5 | 1.1 |
| P21333 | Filamin-A OS=Homo sapiens OX=9606 GN=FLNA<br>PE=1 SV=4 - [FLNA_HUMAN] | FLNA | 26 | 38 | 0.1 | -1.0 | -0.9 | -0.5 | 1.0 | -1.6 |
| P22087 | rRNA 2'-O-methyltransferase fibrillarin OS=Homo<br>sapiens OX=9606 GN=FBRL PE=1 SV=2 - [FBRL_HUMAN] | FBRL | 10 | 15 | 1.1 | -0.3 | 1.7 | -1.0 | -2.1 | -2.0 |
| P22392 | Nucleoside diphosphate kinase B OS=Homo sapiens<br>OX=9606 GN=NME2 PE=1 SV=1 - [NDKB_HUMAN] | NDKB | 6 | 11 | -1.6 | -1.2 | 0.8 | 1.9 | -2.0 | -3.0 |
| P22626 | Heterogeneous nuclear ribonucleoproteins A2/B1<br>OS=Homo sapiens OX=9606 GN=HNRNPA2B1 PE=1<br>SV=2 - [ROA2_HUMAN] | ROA2 | 12 | 23 | 1.0 | -1.0 | 0.8 | -1.9 | -2.1 | -2.7 |
| P23284 | Peptidyl-prolyl cis-trans isomerase B OS=Homo sapiens<br>OX=9606 GN=PPIB PE=1 SV=2 - [PPIB_HUMAN] | PPIB | 3 | 6 | -0.7 | -1.1 | -0.5 | -1.7 | 0.7 | 1.4 |
| P23396 | 40S ribosomal protein S3 OS=Homo sapiens OX=9606<br>GN=RPS3 PE=1 SV=2 - [RS3_HUMAN] | RS3 | 8 | 10 | 0.9 | 0.4 | 2.1 | -1.6 | 0.2 | 0.4 |
| P23526 | Adenosylhomocysteinase OS=Homo sapiens OX=9606<br>GN=AHCY PE=1 SV=4 - [SAHH_HUMAN] | SAHH | 7 | 11 | 0.9 | -0.4 | 1.6 | -1.2 | -1.5 | -1.6 |
| P23528 | Cofilin-1 OS=Homo sapiens OX=9606 GN=CFL1 PE=1<br>SV=3 - [COF1_HUMAN] | COF1 | 3 | 6 | -1.8 | -0.6 | 0.2 | 3.2 | -1.7 | -2.8 |
| P24534 | Elongation factor 1-beta OS=Homo sapiens OX=9606<br>GN=EEF1B2 PE=1 SV=3 - [EF1B_HUMAN] | EF1B | 4 | 6 | 1.9 | -1.1 | 1.2 | -1.7 | -1.3 | -1.7 |
| P25705 | ATP synthase subunit alpha, mitochondrial OS=Homo<br>sapiens OX=9606 GN=ATP5F1A PE=1 SV=1 - [ATPA_HUMAN] | ATPA | 14 | 15 | -0.5 | 0.2 | 2.2 | -0.2 | -1.2 | 0.2 |
| P25786 | Proteasome subunit alpha type-1 OS=Homo sapiens<br>OX=9606 GN=PSMA1 PE=1 SV=1 - [PSA1_HUMAN] | PSA1 | 5 | 7 | 1.4 | -0.7 | 1.3 | -1.9 | -1.2 | -1.3 |
| P25788 | Proteasome subunit alpha type-3 OS=Homo sapiens<br>OX=9606 GN=PSMA3 PE=1 SV=2 - [PSA3_HUMAN] | PSA3 | 5 | 6 | -0.6 | -1.4 | -0.4 | -0.9 | -0.2 | 0.5 |
| P26038 | Moesin OS=Homo sapiens OX=9606 GN=MSN PE=1<br>SV=3 - [MOES_HUMAN] | MOES | 8 | 10 | 0.5 | -2.1 | -1.3 | 1.8 | -1.9 | -0.9 |
| P26373 | 60S ribosomal protein L13 OS=Homo sapiens OX=9606 GN=RPL13 PE=1 SV=4 - [RL13_HUMAN] | RL13 | 11 | 19 | -0.9 | -1.4 | -0.3 | -0.5 | 0.7 | 0.3 |
| P26599 | Polypyrimidine tract-binding protein 1 OS=Homo sapiens OX=9606 GN=PTBP1 PE=1 SV=1 - [PTBP1_HUMAN] | PTBP1 | 8 | 13 | 0.2 | -1.7 | -1.1 | -0.5 | -1.1 | 0.3 |
| P26641 | Elongation factor 1-gamma OS=Homo sapiens OX=9606 GN=EEF1G PE=1 SV=3 - [EF1G_HUMAN] | EF1G | 9 | 11 | 0.2 | -1.6 | -1.0 | -1.2 | -0.6 | 0.9 |
| P27348 | 14-3-3 protein theta OS=Homo sapiens OX=9606 GN=YWHAQ PE=1 SV=1 - [1433T_HUMAN] | 1433T | 6 | 14 | -0.3 | -0.9 | -2.1 | 1.1 | -1.1 | -1.2 |
| P27797 | Calreticulin OS=Homo sapiens OX=9606 GN=CALR PE=1 SV=1 - [CALR_HUMAN] | CALR | 7 | 8 | -0.1 | -2.7 | -1.0 | -1.7 | -0.1 | 0.7 |
| P27824 | Calnexin OS=Homo sapiens OX=9606 GN=CANX PE=1 SV=2 - [CALX_HUMAN] | CALX | 5 | 7 | 1.9 | -1.4 | -2.1 | -0.4 | 1.1 | -1.1 |
| P28066 | Proteasome subunit alpha type-5 OS=Homo sapiens OX=9606 GN=PSMA5 PE=1 SV=3 - [PSA5_HUMAN] | PSA5 | 2 | 3 | -0.2 | -0.9 | -1.2 | 0.3 | -0.2 | -0.1 |
| P28074 | Proteasome subunit beta type-5 OS=Homo sapiens OX=9606 GN=PSMB5 PE=1 SV=3 - [PSB5_HUMAN] | PSB5 | 8 | 12 | -0.9 | -1.6 | -0.4 | 2.1 | -1.6 | -2.1 |
| P28838 | Cytosol aminopeptidase OS=Homo sapiens OX=9606 GN=LAP3 PE=1 SV=3 - [AMPL_HUMAN] | AMPL | 12 | 17 | -0.4 | -1.5 | -0.5 | 1.0 | -0.5 | -1.6 |
| P29401 | Transketolase OS=Homo sapiens OX=9606 GN=TKT PE=1 SV=3 - [TKT_HUMAN] | TKT | 15 | 17 | -0.3 | 0.2 | 2.6 | 1.8 | -1.0 | -3.1 |
| P29692 | Elongation factor 1-delta OS=Homo sapiens OX=9606 GN=EEF1D PE=1 SV=5 - [EF1D_HUMAN] | EF1D | 9 | 9 | -0.2 | -2.8 | -0.6 | -0.8 | -0.8 | -0.7 |
| P30041 | Peroxiredoxin-6 OS=Homo sapiens OX=9606 GN=PRDX6 PE=1 SV=3 - [PRDX6_HUMAN] | PRDX6 | 8 | 11 | -0.7 | -1.1 | -0.1 | -1.5 | 0.1 | 0.6 |
| P30048 | Thioredoxin-dependent peroxide reductase, mitochondrial OS=Homo sapiens OX=9606 GN=PRDX3 PE=1 SV=3 - [PRDX3_HUMAN] | PRDX3 | 3 | 5 | -1.2 | -2.0 | -0.5 | 2.6 | -1.5 | -2.3 |
| P30101 | Protein disulfide-isomerase A3 OS=Homo sapiens OX=9606 GN=PDIA3 PE=1 SV=4 - [PDIA3_HUMAN] | PDIA3 | 9 | 16 | 0.2 | -1.8 | -1.3 | 0.8 | -0.5 | -1.3 |
| P30419 | Glycylpeptide N-tetradecanoyltransferase 1 OS=Homo sapiens OX=9606 GN=NMT1 PE=1 SV=2 - [NMT1_HUMAN] | NMT1 | 5 | 7 | 0.2 | -0.5 | -0.9 | 0.1 | -1.8 | -0.4 |
| P31943 | Heterogeneous nuclear ribonucleoprotein H OS=Homo sapiens OX=9606 GN=HNRNPH1 PE=1 SV=4 - [HNRH1_HUMAN] | HNRH1 | 15 | 29 | -0.6 | -1.5 | -0.2 | -0.7 | -0.9 | 0.6 |
| P31946 | 14-3-3 protein beta/alpha OS=Homo sapiens OX=9606 GN=YWHAB PE=1 SV=3 - [1433B_HUMAN] | 1433B | 8 | 14 | -0.4 | -0.9 | -1.6 | 0.2 | -0.4 | 0.0 |
| P31948 | Stress-induced-phosphoprotein 1 OS=Homo sapiens OX=9606 GN=STIP1 PE=1 SV=1 - [STIP1_HUMAN] | STIP1 | 4 | 5 | 0.1 | -2.1 | -0.9 | -1.2 | -0.7 | 0.8 |
| P32969 | 60S ribosomal protein L9 OS=Homo sapiens OX=9606 GN=RPL9 PE=1 SV=1 - [RL9_HUMAN] | RL9 | 4 | 4 | -0.1 | 0.3 | 1.1 | 1.4 | -1.4 | -1.7 |
| P35232 | Prohibitin OS=Homo sapiens OX=9606 GN=PHB PE=1 SV=1 - [PHB_HUMAN] | PHB | 10 | 11 | -0.5 | -0.8 | -0.3 | -0.3 | 0.4 | 0.1 |
| P35268 | 60S ribosomal protein L22 OS=Homo sapiens OX=9606 GN=RPL22 PE=1 SV=2 - [RL22_HUMAN] | RL22 | 2 | 3 | -0.7 | -1.7 | -0.6 | -2.2 | 1.0 | 2.4 |
| P35270 | Sepiapterin reductase OS=Homo sapiens OX=9606 GN=SPR PE=1 SV=1 - [SPRE_HUMAN] | SPRE | 5 | 6 | -0.2 | -1.0 | -1.4 | -0.8 | -0.9 | 0.0 |
| P35527 | Keratin, type I cytoskeletal 9 OS=Homo sapiens<br>OX=9606 GN=KRT9 PE=1 SV=3 - [K1C9_HUMAN] | K1C9 | 19 | 28 | -2.2 | 0.1 | 0.4 | -0.9 | -1.8 | -0.9 |
| P35579 | Myosin-9 OS=Homo sapiens OX=9606 GN=MYH9<br>PE=1 SV=4 - [MYH9_HUMAN] | MYH9 | 46 | 58 | 1.2 | 0.6 | 1.1 | -0.2 | 0.4 | -0.9 |
| P35908 | Keratin, type II cytoskeletal 2 epidermal OS=Homo<br>sapiens OX=9606 GN=KRT2 PE=1 SV=2 -<br>[K22E_HUMAN] | K22E | 27 | 46 | -2.8 | 0.6 | 0.6 | 1.0 | -1.8 | -2.0 |
| P36578 | 60S ribosomal protein L4 OS=Homo sapiens OX=9606<br>GN=RPL4 PE=1 SV=5 - [RL4_HUMAN] | RL4 | 15 | 16 | -0.2 | -1.4 | -0.6 | -1.3 | -0.5 | 0.3 |
| P36957 | Dihydrolipoyllysine-residue succinyltransferase<br>component of 2-oxoglutarate dehydrogenase complex,<br>mitochondrial OS=Homo sapiens OX=9606 GN=DLST<br>PE=1 SV=4 - [ODO2_HUMAN] | ODO2 | 5 | 6 | 0.2 | -1.5 | -1.1 | 1.5 | -1.8 | -0.5 |
| P37802 | Transgelin-2 OS=Homo sapiens OX=9606<br>GN=TAGLN2 PE=1 SV=3 - [TAGL2_HUMAN] | TAGL2 | 6 | 27 | -0.9 | -1.2 | 0.0 | 1.2 | -1.2 | -1.8 |
| P37837 | Transaldolase OS=Homo sapiens OX=9606<br>GN=TALDO1 PE=1 SV=2 - [TALDO_HUMAN] | TALDO | 10 | 14 | 1.2 | -0.9 | 1.2 | -2.3 | -1.1 | -1.6 |
| P38159 | RNA-binding motif protein, X chromosome OS=Homo<br>sapiens OX=9606 GN=RBMX PE=1 SV=3 -<br>[RBMX_HUMAN] | RBMX | 22 | 26 | 0.1 | 0.4 | 2.1 | -1.5 | -1.9 | -1.7 |
| P38646 | Stress-70 protein, mitochondrial OS=Homo sapiens<br>OX=9606 GN=HSPA9 PE=1 SV=2 -<br>[GRP75_HUMAN] | GRP75 | 14 | 17 | -0.2 | 0.1 | 2.7 | 2.2 | -2.0 | -0.5 |
| P40227 | T-complex protein 1 subunit zeta OS=Homo sapiens<br>OX=9606 GN=CCT6A PE=1 SV=3 - [TCPZ_HUMAN] | TCPZ | 11 | 15 | -0.3 | -1.2 | -0.3 | -0.4 | -0.9 | 0.3 |
| P40925 | Malate dehydrogenase, cytoplasmic OS=Homo sapiens<br>OX=9606 GN=MDH1 PE=1 SV=4 -<br>[MDHC_HUMAN] | MDHC | 4 | 5 | 1.1 | -0.3 | 1.6 | -1.2 | -1.4 | -1.8 |
| P40926 | Malate dehydrogenase, mitochondrial OS=Homo<br>sapiens OX=9606 GN=MDH2 PE=1 SV=3 -<br>[MDHM_HUMAN] | MDHM | 17 | 26 | 1.7 | -0.7 | 1.3 | -2.3 | -1.6 | -1.8 |
| P42330 | Aldo-keto reductase family 1 member C3 OS=Homo<br>sapiens OX=9606 GN=AKR1C3 PE=1 SV=4 -<br>[AK1C3_HUMAN] | AK1C3 | 5 | 5 | -0.4 | -0.1 | -0.4 | -1.5 | -0.1 | 0.6 |
| P43307 | Translocon-associated protein subunit alpha OS=Homo<br>sapiens OX=9606 GN=SSR1 PE=1 SV=3 -<br>[SSRA_HUMAN] | SSRA | 2 | 3 | -0.3 | -1.2 | 0.2 | -0.3 | -0.6 | -0.5 |
| P45880 | Voltage-dependent anion-selective channel protein 2<br>OS=Homo sapiens OX=9606 GN=VDAC2 PE=1 SV=2<br>- [VDAC2_HUMAN] | VDAC2 | 6 | 7 | 1.0 | -1.3 | 0.0 | -2.0 | -1.0 | -1.7 |
| P46777 | 60S ribosomal protein L5 OS=Homo sapiens OX=9606<br>GN=RPL5 PE=1 SV=3 - [RL5_HUMAN] | RL5 | 11 | 12 | 0.4 | -0.1 | 1.3 | -1.0 | -2.1 | -1.7 |
| P46779 | 60S ribosomal protein L28 OS=Homo sapiens OX=9606<br>GN=RPL28 PE=1 SV=3 - [RL28_HUMAN] | RL28 | 6 | 9 | -0.8 | -0.9 | 0.2 | -3.0 | 1.9 | 4.2 |
| P46782 | 40S ribosomal protein S5 OS=Homo sapiens OX=9606<br>GN=RPS5 PE=1 SV=4 - [RS5_HUMAN] | RS5 | 8 | 10 | -0.9 | -0.6 | 0.3 | 1.2 | -1.8 | -2.2 |
| P46783 | 40S ribosomal protein S10 OS=Homo sapiens OX=9606<br>GN=RPS10 PE=1 SV=1 - [RS10_HUMAN] | RS10 | 7 | 10 | -0.7 | -2.0 | -0.6 | -2.2 | 1.0 | 2.6 |
| P47755 | F-actin-capping protein subunit alpha-2 OS=Homo sapiens OX=9606 GN=CAPZA2 PE=1 SV=3 - [CAZA2_HUMAN] | CAZA2 | 4 | 6 | 1.1 | -1.2 | 0.5 | -1.7 | -1.1 | -1.3 |
| P48643 | T-complex protein 1 subunit epsilon OS=Homo sapiens OX=9606 GN=CCT5 PE=1 SV=1 - [TCPE_HUMAN] | TCPE | 5 | 8 | 0.0 | -1.7 | -1.0 | -1.1 | -0.6 | 0.5 |
| P49368 | T-complex protein 1 subunit gamma OS=Homo sapiens OX=9606 GN=CCT3 PE=1 SV=4 - [TCPG_HUMAN] | TCPG | 13 | 24 | 0.2 | -1.4 | -1.0 | -0.2 | -1.3 | 0.0 |
| P49411 | Elongation factor Tu, mitochondrial OS=Homo sapiens OX=9606 GN=TUFM PE=1 SV=2 - [EFTU_HUMAN] | EFTU | 7 | 9 | 1.1 | -0.6 | 1.1 | -0.9 | -0.9 | 0.5 |
| P49773 | Histidine triad nucleotide-binding protein 1 OS=Homo sapiens OX=9606 GN=HINT1 PE=1 SV=2 - [HINT1_HUMAN] | HINT1 | 1 | 1 | -1.4 | -1.7 | -0.5 | 2.3 | -1.4 | -1.5 |
| P50454 | Serpin H1 OS=Homo sapiens OX=9606 GN=SERPINH1 PE=1 SV=2 - [SERPH_HUMAN] | SERPH | 11 | 15 | 0.1 | -1.9 | -1.3 | -1.1 | -0.8 | 0.5 |
| P50990 | T-complex protein 1 subunit theta OS=Homo sapiens OX=9606 GN=CCT8 PE=1 SV=4 - [TCPQ_HUMAN] | TCPQ | 10 | 13 | -0.3 | -1.4 | -0.7 | -1.1 | -0.4 | 0.7 |
| P50991 | T-complex protein 1 subunit delta OS=Homo sapiens OX=9606 GN=CCT4 PE=1 SV=4 - [TCPD_HUMAN] | TCPD | 15 | 18 | -0.7 | -1.3 | -0.7 | 1.1 | -0.7 | -2.1 |
| P50993 | Sodium/potassium-transporting ATPase subunit alpha-2 OS=Homo sapiens OX=9606 GN=ATP1A2 PE=1 SV=1 - [AT1A2_HUMAN] | AT1A2 | 15 | 24 | 0.3 | -1.2 | -0.8 | 1.3 | -2.6 | -1.1 |
| P51148 | Ras-related protein Rab-5C OS=Homo sapiens OX=9606 GN=RAB5C PE=1 SV=2 - [RAB5C_HUMAN] | RAB5C | 7 | 10 | -0.2 | -3.4 | 0.2 | 1.3 | -2.9 | -1.8 |
| P51149 | Ras-related protein Rab-7a OS=Homo sapiens OX=9606 GN=RAB7A PE=1 SV=1 - [RAB7A_HUMAN] | RAB7A | 4 | 7 | -0.5 | -2.0 | -0.7 | 1.1 | -1.0 | -0.8 |
| P51665 | 26S proteasome non-ATPase regulatory subunit 7 OS=Homo sapiens OX=9606 GN=PSMD7 PE=1 SV=2 - [PSMD7_HUMAN] | PSMD7 | 3 | 6 | -0.2 | -1.6 | -1.3 | -1.8 | -1.6 | -3.0 |
| P52272 | Heterogeneous nuclear ribonucleoprotein M OS=Homo sapiens OX=9606 GN=HNRNPM PE=1 SV=3 - [HNRPM_HUMAN] | HNRPM | 39 | 61 | -0.7 | 0.2 | 1.9 | 1.7 | -1.9 | -0.4 |
| P52758 | 2-iminobutanoate/2-iminopropanoate deaminase OS=Homo sapiens OX=9606 GN=RIDA PE=1 SV=1 - [RIDA_HUMAN] | RIDA | 3 | 5 | -0.2 | -1.1 | 0.6 | 1.9 | -1.2 | -1.5 |
| P52895 | Aldo-keto reductase family 1 member C2 OS=Homo sapiens OX=9606 GN=AKR1C2 PE=1 SV=3 - [AK1C2_HUMAN] | AK1C2 | 6 | 9 | 0.0 | -1.1 | -0.9 | 0.5 | -1.2 | 0.0 |
| P52907 | F-actin-capping protein subunit alpha-1 OS=Homo sapiens OX=9606 GN=CAPZA1 PE=1 SV=3 - [CAZA1_HUMAN] | CAZA1 | 4 | 6 | 0.0 | -1.5 | -1.1 | 0.0 | -1.6 | -1.1 |
| P53396 | ATP-citrate synthase OS=Homo sapiens OX=9606 GN=ACLY PE=1 SV=3 - [ACLY_HUMAN] | ACLY | 15 | 21 | 2.0 | -1.2 | -2.2 | -0.9 | -0.1 | 0.4 |
| P55072 | Transitional endoplasmic reticulum ATPase OS=Homo sapiens OX=9606 GN=VCP PE=1 SV=4 - [TERA_HUMAN] | TERA | 21 | 29 | 0.5 | -1.3 | -1.1 | -0.2 | 0.5 | -0.7 |
| P55209 | Nucleosome assembly protein 1-like 1 OS=Homo sapiens OX=9606 GN=NAP1L1 PE=1 SV=1 - [NP1L1_HUMAN] | NP1L1 | 4 | 6 | 0.1 | -2.0 | -1.3 | -2.2 | 0.5 | 1.7 |
| P57735 | Ras-related protein Rab-25 OS=Homo sapiens | RAB25 | 7 | 17 | 0.4 | 0.3 | 0.6 | 1.0 | -1.2 | -1.1 |
|  | OX=9606 GN=RAB25 PE=1 SV=2 - [RAB25_HUMAN] |  |  |  |  |  |  |  |  |  |
| P60174 | Triosephosphate isomerase OS=Homo sapiens OX=9606 GN=TP1I PE=1 SV=3 - [TP1I_HUMAN] | TP1I | 10 | 14 | -0.8 | -1.7 | -0.6 | 3.1 | -1.8 | -2.4 |
| P60709 | Actin, cytoplasmic 1 OS=Homo sapiens OX=9606 GN=ACTB PE=1 SV=1 - [ACTB_HUMAN] | ACTB | 32 | 153 | 1.4 | -1.3 | 0.8 | -2.3 | -0.9 | -0.7 |
| P60842 | Eukaryotic initiation factor 4A-I OS=Homo sapiens OX=9606 GN=EIF4A1 PE=1 SV=1 - [EIF4A1_HUMAN] | EIF4A1 | 14 | 22 | -0.6 | -1.1 | -0.6 | -1.2 | -0.6 | 0.9 |
| P60866 | 40S ribosomal protein S20 OS=Homo sapiens OX=9606 GN=RPS20 PE=1 SV=1 - [RPS20_HUMAN] | RS20 | 4 | 5 | -1.4 | -0.8 | 0.5 | 1.2 | -1.0 | -0.8 |
| P61160 | Actin-related protein 2 OS=Homo sapiens OX=9606 GN=ACTR2 PE=1 SV=1 - [ARP2_HUMAN] | ARP2 | 7 | 7 | 0.2 | -2.3 | -1.7 | -1.1 | -0.5 | 0.5 |
| P61247 | 40S ribosomal protein S3a OS=Homo sapiens OX=9606 GN=RPS3A PE=1 SV=2 - [RS3A_HUMAN] | RS3A | 16 | 22 | 1.6 | -0.5 | 2.2 | -1.3 | -2.2 | -1.9 |
| P61353 | 60S ribosomal protein L27 OS=Homo sapiens OX=9606 GN=RPL27 PE=1 SV=2 - [RL27_HUMAN] | RL27 | 5 | 5 | -0.6 | -2.3 | -0.9 | -4.3 | 2.7 | 3.8 |
| P61604 | 10 kDa heat shock protein, mitochondrial OS=Homo sapiens OX=9606 GN=HSPE1 PE=1 SV=2 - [CH10_HUMAN] | CH10 | 7 | 13 | -1.1 | -1.3 | 0.5 | 1.6 | -1.0 | -0.9 |
| P61978 | Heterogeneous nuclear ribonucleoprotein K OS=Homo sapiens OX=9606 GN=HNRNPK PE=1 SV=1 - [HNRNPK_HUMAN] | HNRNPK | 17 | 28 | 0.2 | -2.1 | -1.5 | -0.5 | -1.0 | 0.4 |
| P61981 | 14-3-3 protein gamma OS=Homo sapiens OX=9606 GN=YWHAG PE=1 SV=2 - [1433G_HUMAN] | 1433G | 6 | 7 | -0.3 | -1.2 | -1.4 | -1.1 | 0.9 | 0.7 |
| P62081 | 40S ribosomal protein S7 OS=Homo sapiens OX=9606 GN=RPS7 PE=1 SV=1 - [RS7_HUMAN] | RS7 | 8 | 13 | -1.4 | -1.3 | 0.1 | 2.0 | -1.3 | -1.7 |
| P62244 | 40S ribosomal protein S15a OS=Homo sapiens OX=9606 GN=RPS15A PE=1 SV=2 - [RS15A_HUMAN] | RS15A | 4 | 4 | -0.4 | -2.8 | -1.2 | -3.2 | 1.7 | 2.7 |
| P62249 | 40S ribosomal protein S16 OS=Homo sapiens OX=9606 GN=RPS16 PE=1 SV=2 - [RS16_HUMAN] | RS16 | 7 | 12 | -0.7 | -1.0 | 0.0 | 0.9 | -1.1 | -0.3 |
| P62258 | 14-3-3 protein epsilon OS=Homo sapiens OX=9606 GN=YWHA E PE=1 SV=1 - [1433E_HUMAN] | 1433E | 9 | 16 | 0.0 | -0.8 | -1.0 | -1.4 | -1.4 | -1.5 |
| P62266 | 40S ribosomal protein S23 OS=Homo sapiens OX=9606 GN=RPS23 PE=1 SV=3 - [RS23_HUMAN] | RS23 | 5 | 9 | -0.8 | -1.0 | 0.4 | 1.2 | -1.4 | -1.9 |
| P62269 | 40S ribosomal protein S18 OS=Homo sapiens OX=9606 GN=RPS18 PE=1 SV=3 - [RS18_HUMAN] | RS18 | 4 | 6 | -0.6 | -1.3 | -0.4 | -2.7 | 1.2 | 2.9 |
| P62277 | 40S ribosomal protein S13 OS=Homo sapiens OX=9606 GN=RPS13 PE=1 SV=2 - [RS13_HUMAN] | RS13 | 8 | 10 | -1.0 | -1.0 | 0.6 | -3.0 | 1.1 | 3.1 |
| P62424 | 60S ribosomal protein L7a OS=Homo sapiens OX=9606 GN=RPL7A PE=1 SV=2 - [RL7A_HUMAN] | RL7A | 6 | 6 | -0.4 | -1.8 | -0.8 | 0.6 | -0.1 | 0.0 |
| P62701 | 40S ribosomal protein S4, X isoform OS=Homo sapiens OX=9606 GN=RPS4X PE=1 SV=2 - [RS4X_HUMAN] | RS4X | 4 | 4 | -0.1 | -1.1 | -1.1 | 2.4 | -1.9 | -3.1 |
| P62805 | Histone H4 OS=Homo sapiens OX=9606 GN=HIST1H4A PE=1 SV=2 - [H4_HUMAN] | H4 | 11 | 47 | -1.6 | -1.0 | 0.5 | 2.8 | -3.1 | -3.8 |
| P62826 | GTP-binding nuclear protein Ran OS=Homo sapiens OX=9606 GN=RAN PE=1 SV=3 - [RAN_HUMAN] | RAN | 8 | 12 | -1.4 | -1.5 | -0.2 | 2.9 | -2.0 | -2.5 |
| P62851 | 40S ribosomal protein S25 OS=Homo sapiens OX=9606 GN=RPS25 PE=1 SV=1 - [RS25_HUMAN] | RS25 | 4 | 6 | -1.6 | -1.0 | 0.7 | 2.9 | -2.1 | -2.2 |
| P62857 | 40S ribosomal protein S28 OS=Homo sapiens OX=9606 GN=RPS28 PE=1 SV=1 - [RS28_HUMAN] | RS28 | 5 | 14 | -2.8 | -1.5 | 0.3 | -3.1 | 2.0 | 3.9 |
| P62899 | 60S ribosomal protein L31 OS=Homo sapiens OX=9606 GN=RPL31 PE=1 SV=1 - [RL31_HUMAN] | RL31 | 3 | 4 | -0.8 | -1.5 | -0.4 | -2.8 | 1.1 | 2.5 |
| P62910 | 60S ribosomal protein L32 OS=Homo sapiens OX=9606 GN=RPL32 PE=1 SV=2 - [RL32_HUMAN] | RL32 | 4 | 5 | -0.9 | -0.9 | 0.9 | 1.3 | -1.1 | -1.8 |
| P62917 | 60S ribosomal protein L8 OS=Homo sapiens OX=9606 GN=RPL8 PE=1 SV=2 - [RL8_HUMAN] | RL8 | 12 | 13 | 1.0 | -0.5 | 1.6 | -1.1 | -1.5 | -1.8 |
| P62937 | Peptidyl-prolyl cis-trans isomerase A OS=Homo sapiens OX=9606 GN=PPIA PE=1 SV=2 - [PPIA_HUMAN] | PPIA | 10 | 22 | -1.4 | -1.3 | 0.4 | 1.9 | -1.5 | -2.7 |
| P62979 | Ubiquitin-40S ribosomal protein S27a OS=Homo sapiens OX=9606 GN=RPS27A PE=1 SV=2 - [RS27A_HUMAN] | RS27A | 8 | 19 | -1.5 | -1.7 | 0.5 | 2.2 | -2.3 | -2.7 |
| P62995 | Transformer-2 protein homolog beta OS=Homo sapiens OX=9606 GN=TRA2B PE=1 SV=1 - [TRA2B_HUMAN] | TRA2B | 11 | 16 | 0.9 | -1.1 | 0.9 | 0.2 | -1.9 | -0.8 |
| P63104 | 14-3-3 protein zeta/delta OS=Homo sapiens OX=9606 GN=YWHAZ PE=1 SV=1 - [1433Z_HUMAN] | 1433Z | 14 | 36 | -0.9 | -1.7 | -0.3 | -0.4 | 0.2 | 0.3 |
| P63244 | Receptor of activated protein C kinase 1 OS=Homo sapiens OX=9606 GN=RACK1 PE=1 SV=3 - [RACK1_HUMAN] | RACK1 | 13 | 19 | -1.1 | -1.2 | -0.4 | -2.2 | -2.2 | -1.3 |
| P68104 | Elongation factor 1-alpha 1 OS=Homo sapiens OX=9606 GN=EEF1A1 PE=1 SV=1 - [EF1A1_HUMAN] | EF1A1 | 16 | 31 | -0.9 | -1.7 | -1.0 | 1.3 | -0.9 | -2.5 |
| P68133 | Actin, alpha skeletal muscle OS=Homo sapiens OX=9606 GN=ACTA1 PE=1 SV=1 - [ACTS_HUMAN] | ACTS | 16 | 61 | 0.9 | -1.7 | -2.3 | -2.4 | -2.1 | -1.4 |
| P68371 | Tubulin beta-4B chain OS=Homo sapiens OX=9606 GN=TUBB4B PE=1 SV=1 - [TBB4B_HUMAN] | TBB4B | 20 | 44 | -0.3 | -1.3 | -0.4 | 0.9 | -0.6 | -1.2 |
| P68431 | Histone H3.1 OS=Homo sapiens OX=9606 GN=HIST1H3A PE=1 SV=2 - [H31_HUMAN] | H31 | 8 | 13 | 0.2 | 1.2 | 1.7 | 2.5 | -1.9 | -2.4 |
| P78371 | T-complex protein 1 subunit beta OS=Homo sapiens OX=9606 GN=CCT2 PE=1 SV=4 - [TCPB_HUMAN] | TCPB | 17 | 19 | -0.5 | -1.6 | -0.9 | 1.8 | -0.3 | -2.4 |
| P81605 | Dermcidin OS=Homo sapiens OX=9606 GN=DCD PE=1 SV=2 - [DCD_HUMAN] | DCD | 4 | 5 | -1.4 | 0.3 | 0.7 | 1.6 | -2.4 | -2.7 |
| Q00534 | Cyclin-dependent kinase 6 OS=Homo sapiens OX=9606 GN=CDK6 PE=1 SV=1 - [CDK6_HUMAN] | CDK6 | 8 | 11 | -0.1 | -0.9 | -1.1 | -1.1 | -1.0 | -1.3 |
| Q00610 | Clathrin heavy chain 1 OS=Homo sapiens OX=9606 GN=CLTC PE=1 SV=5 - [CLH1_HUMAN] | CLH1 | 26 | 29 | 0.3 | -1.4 | -1.2 | -2.1 | 1.0 | 2.8 |
| Q00839 | Heterogeneous nuclear ribonucleoprotein U OS=Homo sapiens OX=9606 GN=HNRNPU PE=1 SV=6 - [HNRPU_HUMAN] | HNRPU | 14 | 15 | 0.0 | -1.0 | -0.7 | 1.3 | -2.8 | -1.0 |
| Q02878 | 60S ribosomal protein L6 OS=Homo sapiens OX=9606 GN=RPL6 PE=1 SV=3 - [RL6_HUMAN] | RL6 | 5 | 7 | -0.2 | -0.9 | -1.9 | -3.5 | 2.0 | 1.3 |
| Q03252 | Lamin-B2 OS=Homo sapiens OX=9606 GN=LMNB2 PE=1 SV=4 - [LMNB2_HUMAN] | LMNB2 | 21 | 26 | -0.2 | -1.7 | -0.8 | 2.1 | -0.5 | -2.6 |
| Q04828 | Aldo-keto reductase family 1 member C1 OS=Homo sapiens OX=9606 GN=AKR1C1 PE=1 SV=1 - [AK1C1_HUMAN] | AK1C1 | 5 | 7 | -0.1 | -1.1 | -0.8 | -1.3 | -1.2 | -1.0 |
| Q04917 | 14-3-3 protein eta OS=Homo sapiens OX=9606 GN=YWHAH PE=1 SV=4 - [1433F_HUMAN] | 1433F | 6 | 6 | -1.5 | -0.3 | -1.7 | -0.7 | -0.6 | 0.2 |
| Q06830 | Peroxisredoxin-1 OS=Homo sapiens OX=9606 GN=PRDX1 PE=1 SV=1 - [PRDX1_HUMAN] | PRDX1 | 12 | 30 | -1.3 | -1.3 | -0.2 | 1.4 | -1.7 | -1.9 |
| Q07065 | Cytoskeleton-associated protein 4 OS=Homo sapiens OX=9606 GN=CKAP4 PE=1 SV=2 - [CKAP4_HUMAN] | CKAP4 | 5 | 6 | 0.1 | -2.4 | -1.1 | -1.7 | 0.3 | 1.4 |
| Q07955 | Serine/arginine-rich splicing factor 1 OS=Homo sapiens OX=9606 GN=SRSF1 PE=1 SV=2 - [SRSF1_HUMAN] | SRSF1 | 15 | 22 | -0.7 | -1.6 | -0.7 | -1.5 | -1.0 | -1.4 |
| Q08257 | Quinone oxidoreductase OS=Homo sapiens OX=9606 GN=CRYZ PE=1 SV=1 - [QOR_HUMAN] | QOR | 7 | 13 | -0.3 | -1.1 | -1.8 | -1.4 | -1.0 | -1.0 |
| Q13162 | Peroxisredoxin-4 OS=Homo sapiens OX=9606 GN=PRDX4 PE=1 SV=1 - [PRDX4_HUMAN] | PRDX4 | 5 | 12 | -0.8 | -0.6 | -0.6 | 1.2 | -1.2 | -1.4 |
| Q13442 | 28 kDa heat- and acid-stable phosphoprotein OS=Homo sapiens OX=9606 GN=PDAP1 PE=1 SV=1 - [HAP28_HUMAN] | HAP28 | 4 | 4 | -0.5 | 0.2 | 0.6 | 0.7 | -1.1 | -0.7 |
| Q14315 | Filamin-C OS=Homo sapiens OX=9606 GN=FLNC PE=1 SV=3 - [FLNC_HUMAN] | FLNC | 34 | 45 | -1.5 | 0.1 | 0.6 | 1.6 | -2.8 | -1.3 |
| Q14376 | UDP-glucose 4-epimerase OS=Homo sapiens OX=9606 GN=GALE PE=1 SV=2 - [GALE_HUMAN] | GALE | 2 | 2 | 0.0 | -1.6 | -1.0 | 0.0 | -0.9 | -0.1 |
| Q14847 | LIM and SH3 domain protein 1 OS=Homo sapiens OX=9606 GN=LASP1 PE=1 SV=2 - [LASP1_HUMAN] | LASP1 | 8 | 8 | 0.2 | -1.6 | -1.5 | 1.3 | -0.4 | -1.6 |
| Q14974 | Importin subunit beta-1 OS=Homo sapiens OX=9606 GN=KPNB1 PE=1 SV=2 - [IMB1_HUMAN] | IMB1 | 10 | 13 | -1.4 | -0.5 | 0.4 | -0.1 | 0.3 | -0.2 |
| Q15008 | 26S proteasome non-ATPase regulatory subunit 6 OS=Homo sapiens OX=9606 GN=PSMD6 PE=1 SV=1 - [PSMD6_HUMAN] | PSMD6 | 8 | 15 | -0.1 | -1.6 | -0.7 | -0.7 | -1.5 | -1.4 |
| Q15084 | Protein disulfide-isomerase A6 OS=Homo sapiens OX=9606 GN=PDIA6 PE=1 SV=1 - [PDIA6_HUMAN] | PDIA6 | 10 | 16 | -0.4 | -0.1 | 1.5 | -2.4 | 0.5 | 1.2 |
| Q15181 | Inorganic pyrophosphatase OS=Homo sapiens OX=9606 GN=PPA1 PE=1 SV=2 - [IPYR_HUMAN] | IPYR | 5 | 8 | 1.2 | -0.8 | 1.3 | -1.2 | -2.0 | -2.3 |
| Q15233 | Non-POU domain-containing octamer-binding protein OS=Homo sapiens OX=9606 GN=NONO PE=1 SV=4 - [NONO_HUMAN] | NONO | 11 | 14 | 0.0 | -0.3 | 2.6 | -0.8 | -1.0 | 0.2 |
| Q15366 | Poly(rC)-binding protein 2 OS=Homo sapiens OX=9606 GN=PCBP2 PE=1 SV=1 - [PCBP2_HUMAN] | PCBP2 | 7 | 9 | -0.5 | -1.4 | -1.3 | -2.7 | -1.3 | -1.2 |
| Q15417 | Calponin-3 OS=Homo sapiens OX=9606 GN=CNN3 PE=1 SV=1 - [CNN3_HUMAN] | CNN3 | 9 | 11 | 0.7 | -0.5 | 1.2 | -1.4 | -1.2 | -1.3 |
| Q5VTE0 | Putative elongation factor 1-alpha-like 3 OS=Homo sapiens OX=9606 GN=EEF1A1P5 PE=5 SV=1 - [EF1A3_HUMAN] | EF1A3 | 14 | 26 | 0.1 | -2.6 | -0.8 | -1.9 | -0.3 | 1.1 |
| Q6PI78 | Transmembrane protein 65 OS=Homo sapiens OX=9606 GN=TMEM65 PE=1 SV=2 - [TMM65_HUMAN] | TMM65 | 1 | 1 | 0.2 | -1.6 | -1.2 | -0.3 | -1.1 | 0.2 |
| Q6ZRQ5 | Protein MMS22-like OS=Homo sapiens OX=9606 | MMS22 | 15 | 25 | 1.2 | -0.3 | 3.1 | -0.1 | 0.1 | 0.0 |
|  | GN=MMS22L PE=1 SV=3 - [MMS22_HUMAN] |  |  |  |  |  |  |  |  |  |
| Q86XK2 | F-box only protein 11 OS=Homo sapiens OX=9606 GN=FBXO11 PE=1 SV=3 - [FBX11_HUMAN] | FBX11 | 8 | 13 | 0.9 | -3.0 | -0.6 | -0.1 | 0.1 | 0.0 |
| Q8IYB7 | DIS3-like exonuclease 2 OS=Homo sapiens OX=9606 GN=DIS3L2 PE=1 SV=4 - [DI3L2_HUMAN] | DI3L2 | 7 | 9 | -0.6 | -1.2 | -0.3 | 0.3 | -3.9 | 0.7 |
| Q8N257 | Histone H2B type 3-B OS=Homo sapiens OX=9606 GN=HIST3H2BB PE=1 SV=3 - [H2B3B_HUMAN] | H2B3B | 10 | 49 | -1.2 | -0.8 | 0.4 | 1.0 | -1.0 | -1.7 |
| Q8NBS9 | Thioredoxin domain-containing protein 5 OS=Homo sapiens OX=9606 GN=TXNDC5 PE=1 SV=2 - [TXND5_HUMAN] | TXND5 | 5 | 7 | 0.1 | -1.5 | -1.1 | -0.9 | -1.0 | 0.4 |
| Q969Z0 | FAST kinase domain-containing protein 4 OS=Homo sapiens OX=9606 GN=TBG4 PE=1 SV=1 - [FAKD4_HUMAN] | FAKD4 | 10 | 13 | 0.0 | 0.2 | 1.8 | -0.1 | -0.2 | -0.2 |
| Q96C19 | EF-hand domain-containing protein D2 OS=Homo sapiens OX=9606 GN=EFHD2 PE=1 SV=1 - [EFHD2_HUMAN] | EFHD2 | 7 | 8 | -0.1 | -1.0 | -1.1 | -1.5 | -0.9 | -0.8 |
| Q99497 | Protein/nucleic acid deglycase DJ-1 OS=Homo sapiens OX=9606 GN=PARK7 PE=1 SV=2 - [PARK7_HUMAN] | PARK7 | 4 | 14 | -0.6 | -0.9 | -0.1 | 2.1 | -1.4 | -1.6 |
| Q99623 | Prohibitin-2 OS=Homo sapiens OX=9606 GN=PHB2 PE=1 SV=2 - [PHB2_HUMAN] | PHB2 | 11 | 16 | -0.4 | -2.0 | -0.5 | -1.7 | -1.3 | -1.2 |
| Q99714 | 3-hydroxyacyl-CoA dehydrogenase type-2 OS=Homo sapiens OX=9606 GN=HSD17B10 PE=1 SV=3 - [HCD2_HUMAN] | HCD2 | 5 | 7 | -0.3 | -1.3 | -0.8 | 0.8 | -1.0 | -0.9 |
| Q99729 | Heterogeneous nuclear ribonucleoprotein A/B OS=Homo sapiens OX=9606 GN=HNRNPAB PE=1 SV=2 - [ROAA_HUMAN] | ROAA | 4 | 4 | 0.2 | -1.9 | -0.9 | 0.0 | -1.3 | 0.2 |
| Q99832 | T-complex protein 1 subunit eta OS=Homo sapiens OX=9606 GN=CCT7 PE=1 SV=2 - [TCPH_HUMAN] | TCPH | 11 | 15 | 0.1 | -1.3 | -0.6 | -0.6 | -1.0 | 0.2 |
| Q9BPU6 | Dihydropyrimidinase-related protein 5 OS=Homo sapiens OX=9606 GN=DPYSL5 PE=1 SV=1 - [DPYL5_HUMAN] | DPYL5 | 3 | 6 | -0.8 | -0.9 | 0.4 | 1.4 | -1.2 | -1.8 |
| Q9BQE3 | Tubulin alpha-1C chain OS=Homo sapiens OX=9606 GN=TUBA1C PE=1 SV=1 - [TBA1C_HUMAN] | TBA1C | 21 | 41 | -0.5 | -1.5 | -0.5 | -0.6 | -1.2 | 0.4 |
| Q9BTM1 | Histone H2A.J OS=Homo sapiens OX=9606 GN=H2AFJ PE=1 SV=1 - [H2AJ_HUMAN] | H2AJ | 6 | 16 | -1.4 | -1.2 | -0.2 | 1.1 | -1.1 | -1.7 |
| Q9H0C2 | ADP/ATP translocase 4 OS=Homo sapiens OX=9606 GN=SLC25A31 PE=2 SV=1 - [ADT4_HUMAN] | ADT4 | 10 | 12 | 0.2 | 0.3 | 1.6 | -0.7 | 1.0 | -0.5 |
| Q9H0E2 | Toll-interacting protein OS=Homo sapiens OX=9606 GN=TOLLIP PE=1 SV=1 - [TOLIP_HUMAN] | TOLIP | 3 | 3 | -0.2 | -0.4 | 1.2 | 0.6 | -0.7 | -0.2 |
| Q9NR45 | Sialic acid synthase OS=Homo sapiens OX=9606 GN=NANS PE=1 SV=2 - [SIAS_HUMAN] | SIAS | 8 | 10 | 1.0 | 0.1 | 1.6 | -1.5 | -0.9 | -0.9 |
| Q9NRX3 | NADH dehydrogenase [ubiquinone] 1 alpha subcomplex subunit 4-like 2 OS=Homo sapiens OX=9606 GN=NDUFA4L2 PE=3 SV=1 - [NUA4L_HUMAN] | NDUFA4L2 | 2 | 2 | -0.1 | 0.1 | 0.0 | 0.4 | -1.7 | -0.9 |
| Q9NSD9 | Phenylalanine--tRNA ligase beta subunit OS=Homo sapiens OX=9606 GN=FARSB PE=1 SV=3 - [SYFB_HUMAN] | SYFB | 4 | 4 | 0.3 | -1.4 | -1.1 | 0.2 | -1.7 | -0.2 |
| Q9P219 | Protein Daple OS=Homo sapiens OX=9606 GN=CCDC88C PE=1 SV=3 - [DAPLE_HUMAN] | DAPLE | 36 | 47 | -0.1 | 0.8 | 0.8 | 0.1 | -1.5 | -0.2 |
| Q9UBQ5 | Eukaryotic translation initiation factor 3 subunit K OS=Homo sapiens OX=9606 GN=EIF3K PE=1 SV=1 - [EIF3K_HUMAN] | EIF3K | 3 | 7 | -0.7 | -0.8 | -0.2 | 0.1 | -1.4 | 0.6 |
| Q9UHV9 | Prefoldin subunit 2 OS=Homo sapiens OX=9606 GN=PFDN2 PE=1 SV=1 - [PFD2_HUMAN] | PFD2 | 3 | 4 | 0.2 | -0.3 | 0.3 | 0.5 | -0.8 | -0.4 |
| Q9UL46 | Proteasome activator complex subunit 2 OS=Homo sapiens OX=9606 GN=PSME2 PE=1 SV=4 - [PSME2_HUMAN] | PSME2 | 7 | 14 | 1.0 | -0.9 | 0.9 | -1.4 | -1.3 | -1.1 |
| Q9ULE4 | Protein FAM184B OS=Homo sapiens OX=9606 GN=FAM184B PE=2 SV=3 - [F184B_HUMAN] | F184B | 18 | 20 | 1.9 | -0.8 | 1.2 | -1.9 | -2.0 | -2.0 |
| Q9UMS4 | Pre-mRNA-processing factor 19 OS=Homo sapiens OX=9606 GN=PRPF19 PE=1 SV=1 - [PRP19_HUMAN] | PRP19 | 2 | 3 | 0.2 | -1.3 | -1.2 | -0.3 | -1.5 | -0.1 |
| Q9UNM6 | 26S proteasome non-ATPase regulatory subunit 13 OS=Homo sapiens OX=9606 GN=PSMD13 PE=1 SV=2 - [PSD13_HUMAN] | PSD13 | 1 | 1 | -0.2 | -1.2 | -0.9 | -1.8 | -1.5 | -2.3 |
| Q9UQ05 | Potassium voltage-gated channel subfamily H member 4 OS=Homo sapiens OX=9606 GN=KCNH4 PE=2 SV=1 - [KCNH4_HUMAN] | KCNH4 | 11 | 13 | -0.2 | -4.0 | -6.6 | 0.9 | -2.6 | -0.9 |
| Q9Y230 | RuvB-like 2 OS=Homo sapiens OX=9606 GN=RUVBL2 PE=1 SV=3 - [RUVB2_HUMAN] | RUVB2 | 6 | 8 | 0.2 | -1.6 | -1.3 | -0.5 | -1.1 | 0.2 |
| Q9Y265 | RuvB-like 1 OS=Homo sapiens OX=9606 GN=RUVBL1 PE=1 SV=1 - [RUVB1_HUMAN] | RUVB1 | 3 | 5 | 0.2 | -1.4 | -1.2 | 2.7 | -3.3 | -0.3 |
| Q9Y617 | Phosphoserine aminotransferase OS=Homo sapiens OX=9606 GN=PSAT1 PE=1 SV=2 - [SERC_HUMAN] | SERC | 1 | 2 | -0.6 | -1.2 | -0.9 | -2.0 | -1.5 | -1.1 |
| Q2TB90 | Putative hexokinase HKDC1 OS=Homo sapiens OX=9606 GN=HKDC1 PE=1 SV=3 - [HKDC1_HUMAN] | HKDC1 | 14 | 18 | 1.0 | 0.1 | 1.5 | 1.7 | -3.1 | -1.3 |
| Q86YZ3 | Hornerin OS=Homo sapiens OX=9606 GN=HRNR PE=1 SV=2 - [HORN_HUMAN] | HORN | 21 | 31 | -1.5 | -0.2 | -1.1 | 1.4 | -1.9 | -1.8 |

**Figure 5:**
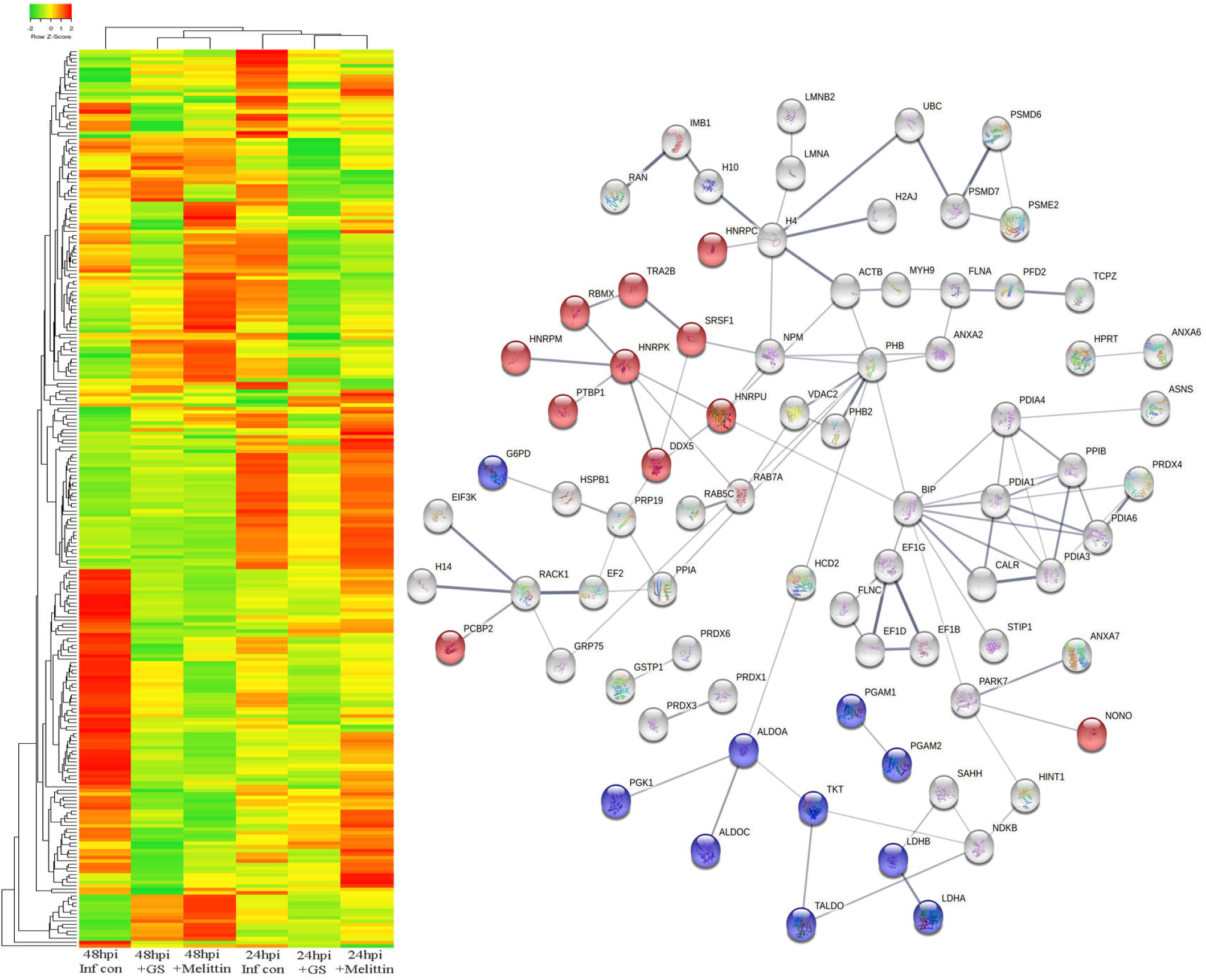
a. Heat map expression of proteins in Vero cells post infection and peptide treatment. b. Network pathway analysis of the differentially expressed protein based on STRING analysis.

A total of 125 proteins were selected for the Network and pathway analysis involving STRING v11.5. Based on Gene ontological functional enrichment analysis it was found that cellular process, biological regulation and regulation of biological process are the most highly associated biological process; binding, proteins binding and heterocyclic compound binding are the most associated molecular functions and cellular anatomical entity, intracellular and organelle are the most occurred cellular components. Carbon metabolism (Blue color nodes), pentose phosphate pathway (Blue color nodes) and mRNA processing (Red color nodes) were most prominent local network cluster associated with STRING analysis (Figure 5b).

### Interaction of gramicidin S and melittin with RBD domain of spike protein: *In silico* analysis

Both gramicidin S and melittin are membrane-active peptides and exert their antimicrobial activity by interacting with membrane components (11, 12). Both the peptides may exert their antiviral activity by targeting multiple regions of the virus. The ability of the peptides to bind to the receptor binding domain (RBD) of the SARS-CoV-2 spike protein was examined by molecular docking using web version of the program ZDOCK (21). The structures shown in Figure 6 show that both gramicidin S and melittin can bind to the RBD binding domain. Panel A shows the crystal structure of RBD-ACE2 complex. The interface between RBD and ACE are shown in violet color and stick representation respectively. The models of gramicidin S and melittin are shown in panels B and C respectively. The LigPlot [22] of the interacting amino acids are shown in panels D for gramicidin S and E for melittin. The residues highlighted in yellow are involved in RBD binding to ACE2 [8]. The modeling study indicates that both the peptides can bind to RBD although their sequences are considerably different from the RBD binding region of ACE2.

**Figure 6:**
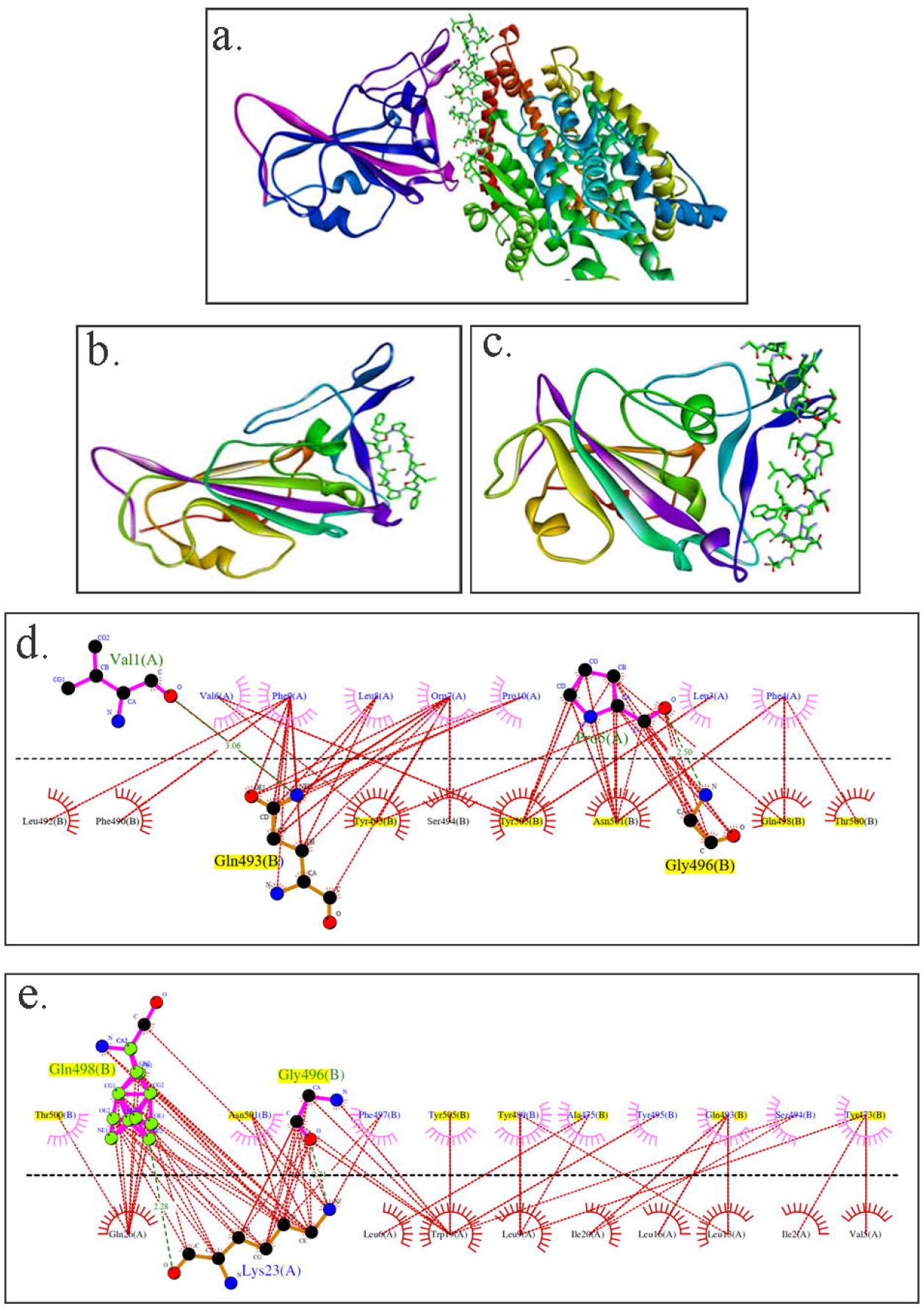
Structures of RBD of spike proteins and peptides and RBD binding region of ACE 2.(a) RBD and ACE, (b) RBD and gramicidin S, (c)RBD and melittin. The ACE2 binding region in RBD is colored violet. In ACE2, gramicidin S and melittin, the peptide regions binding to RBD are represented as sticks. LigPlots of interaction between ACE2 binding domain of the spike protein and (d) gramicidin S and (e) melittin. The residues of the RBD involved in binding to ACE2 are in yellow.

## Discussion

During the past two decades, the world had witnessed infection by three highly pathogenic human corona viruses namely, SARS-Co-V, MERS, SARS-CoV-2 [23,24]. They belong to the group β-coronavirus and have the ability to cross animal-human barriers and cause serious illness in humans. The timely development of specific antivirals is of utmost importance. The development of vaccines at “warp speed” has led to decrease in mortality and serious illness caused by SARS-CoV-2[2]. However, it is still not well established that whether vaccines are equally effective against the several variants that are emerging or the time frame when immunity will be present. To-date, there is no specific drug for SARS-CoV-2. Hence, there is clearly an urgent need to develop therapeutic molecules that would effectively neutralize the virus rather than repurposing drugs used to treat other viral infections. In case of HIV, drugs specific to the virus has been effective to treat the disease although there is still no effective vaccine. Likewise, malaria can be treated with specific drugs rather than vaccines.

However, development of specific therapeutic antiviral drugs for clinical use in a short span is extremely challenging. Repurposing of drugs already known for their therapeutic effects have been extensively screened and tested for inhibition of SARS-CoV-2 [1, 5]. While many repurposed drugs have shown excellent anti SARS-CoV-2 activity *in vitro*, they have had very little success when used clinically. We have shown that host defence peptide such as β-defensin may have a role in infection by SARS-CoV-2 as they are down regulated in COVID-19 patients [25].

Bee venom (BV) has immunosuppressive activity and is generally used in contemporary medicine to treat Multiple Sclerosis, Parkinson’s disease, and arthritis. Bee venom activates foxP3-expressing cells, CD25 and CD4+ T cells, and thus modulates the IgE antibody ratio, resulting in a variety of allergic reactions to antigens [26]. This immunosuppressive activity was observed in Wuhan beekeepers against COVID-19 [26]. Melittin is the main component of bee venom, and it active against both enveloped and non-enveloped viruses by activating the Toll-like receptors (TLRs) pathway, which reduces inflammatory cytokines like nuclear factor-kappa B (NF-kB), extracellular signal-regulated kinases (ERK1/2), and protein kinase Akt [26]. Melittin exhibited antiviral activity against several viruses *in vitro* [16]. Significant antiviral activity of the sitagliptin and melittin-nanoconjugates complex (IC50 value-8.439μM) was observed against SARS-CoV-2 *in vitro* [17]. Binding of sitagliptin and melittin nano conjugates to the active site of SARS-CoV-2 3CLpro (a protease) was also observed through molecular docking [17].

Gramicidin S has potent antibacterial and fungicidal activity [11]. Molecular docking revealed that Gramicidin S has a binding affinity of 11.4 kcal/mol to the SARS-CoV-2 spike glycoprotein and SARS-CoV-2 papain like protease, implying that gramicidin S could be an effective drug against the SARS-CoV-2 virus [27].

SARS-Cov-2 is an enveloped virus, with the viral membrane essential for its integrity and function [1, 8]. We reasoned that membrane-active peptides would disrupt the viral membrane and render the virus ineffective. We have investigated the antiviral activity of two well studied membrane-active antibacterial peptides gramicidin S and melittin.

Our *in vitro* studies using gramicidin S and melittin showed EC_50_ value of 1.571μg for gramicidin S and 0.656μg for melittin (Figure 1). The results were comparable with remdesivir which we used as the assay controlin our study (Figure 5). Our immunofluorescence studies are in agreement with our RT-qPCR data showing the decrease in the viral load in the gramicidin S and melittin treated experimental groups compared with peptide non treated group (Figure 4). Molecular docking studies indicate that both the peptides can bind to the RBD of the spike protein (Figure 6). It is also possible that the peptides bind to RBD of the spike protein and prevent interaction with ACE2 receptor. In fact, membrane-active antibacterial peptides have multiple targets in bacteria. It is conceivable that gramicidin S and melittin also have multiple targets on the virus thereby acting as an effective antiviral agent.

Proteomics studies indicate that metabolic change caused by SARS-CoV-2 pathogenesis result in long-term metabolic disorders in COVID-19 patients, and this varies according to pathogen severity. Carbon sources, specifically glycolysis and glutamino lysis pathways, have been found to play critical roles in SARS-CoV-2 viral replication and production [28]. Non-oxidative pentose phosphate pathways (PPP) are also involved in viral replication; Transketolase is a key mediator enzyme of PPP involved in ribonucleotide production. Benfooxythiamine, a TKT inhibitor, acted against SARS-CoV-2 infection and inhibited the viral replication [29]. Our findings from proteomic pathway analysis show that several proteins are strongly associated with carbon metabolism and non-oxidative PPP. Specifically, LDHA, LDHB, ALDOA, TALDO, PGK1, and PGAM2 proteins were found to be upregulated in early viral replication, i.e. at 24hr viral induced cell control, and vice versa in gramicidin-treated cells. TKT was found to be up regulated after 48 hours of viral induced cell control, but it was substantially down regulated by melittin treatment. It suggests that gramicidin and melittin may function as viral inhibitors by suppressing intercellular metabolic regulators.

Another prominent pathway, mRNA processing, was identified during the local network pathway analysis. According to a recent study on SARS-CoV-2 RNA host protein interaction, the majority of virus induced host RNA binding proteins prevent the virus induced cell death. Several mRNA binding proteins, including Heterogeneous nuclear ribonucleoproteins (HNRNPs), dead box RNA helicases (DDX), and NONO, were activated during the innate immune response to SARS-CoV-2 infection [30]. In our study, NONO, DDX5, RBMX, and HNRPM proteins were found to be up regulated with more than 1 log fold change in melittin-treated SARS-CoV-2 infected cells. This finding suggests that melittin may have antiviral activity during the early stages of viral infection by activating host RNA binding proteins.

The antimicrobial activity of gramicidin S and melittin have been well characterized. Our results strongly argue for development of peptides, gramicidin S and Melittin, as potent therapeutic candidates to treat SARS-CoV-2 and possibly other “cold” causing viruses which are also enveloped viruses, for which there are no effective vaccines.

## Materials and Methods

### Peptides

Gramicidin S and melittin were procured from commercial sources (Gramicidin S: 368108, Calbiochem CA, USA Melittin: M4171 from Sigma Chemical Co, USA). They were characterized by HPLC and mass spectrometry and found to be > 95% pure.

### Cell viability using MTT assay

The Vero cells were plated in 96 well culture plate and incubated at 37°C with 5% CO_2_. After attaining 90-95% cell confluency, different concentrations of gramicidin S and melittin (0.5, 0.7, 3, 5 μg for both) were added to the cells to check the effect of the peptides on the cells for 24 hours. After 24 hours, 100 μl (50 μg) of MTT substrate was added to the cells and the plate was continued to incubate for 3 hours at 37°C with 5% CO_2_. Later the formazan crystals formed were dissolved in 100 μl of DMSO and the absorbance was measured at 570 nm in Multimode Micro plate reader (Synergy HIM).

### RT-qPCR assay

The effect of gramicidin S and melittin was tested against the SARS-CoV-2 with different concentrations. Remdesivir was run as an assay control. The titers for the virus were adjusted such that there was only viral replication and no cytolysis. Briefly, the virus (MOI 0.1) was pre-incubated with different concentrations of melittin and gramicidin S (0.1 – 10 μg) for an hour at 37°C. After the incubation, virus inoculum containing melittin & gramicidin S was added to the Vero cells in duplicates (50μl /well). Remdesivir (1μM) was added to the Vero cells without pre-incubation as in the case of peptides. All the experimental groups were left for infection for 3 hours while maintaining at 37°C with 5% CO_2_. Postinfection (PI), media containing viral inoculum and the gramicidin S and melittin was removed and replaced with 200μl of fresh DMEM media containing 10% FBS and the experimental groups were maintained for varying time points in an incubator maintained at 37°C with 5% CO_2_. Post-incubation, cell supernatants from the experimental groups were collected and spun for 10 min at 6,000 g to remove debris and the supernatant was transferred to fresh collection tubes and later were processed to isolate viral RNA. RNA was extracted from 200 μL aliquots of sample supernatant using the MagMAX™ Viral/Pathogen Extraction Kit (Applied Biosystems, Thermofisher). Extraction of viral RNA was carried out according to the manufacturer’s instructions. Briefly, the viral supernatants from the test groups were added into the deep well plate (KingFisher™Thermo Scientific) along with a lysis buffer containing the following components - 260 μL, MagMAX™ Viral/Pathogen Binding Solution; 10 μL, MVP-II Binding Beads; 5 μL, MagMAX™Viral /Pathogen Proteinase-K, for 200μL of sample. RNA extraction was performed using KingFisher Flex (version 1.01, Thermo Scientific) according to manufactures instructions. The eluted RNA was immediately stored in −80 □C until further use.

The detection of SARS-CoV-2 was done using COVID-19 RT-PCR Detection Kit (Fosun 2019-nCoV qPCR, Shanghai Fosun Long March Medical Science Co. Ltd.) according to the manufacturer’s instructions. The kit detects Envelope gene (E; ROX labelled), Nucleocapsid gene (N-JOE labelled) and Open Reading Frame1ab (ORF1ab, FAM labelled) specific to SARS-CoV-2 for detection and amplification of the cDNA. SARS-CoV-2 cDNA (Ct~28) was used as a positive control. The log viral particles and a semi–log graph was plotted through the linear regression equation obtained using the RNA extracted from the known viral particles by RT-qPCR, using N-gene specific to SARS CoV-2 virus.

### Imunocytochemistry

The Vero cells were seeded in 6-well plate with the sterile glass cover slips. Cells at 90-95 % confluency were considered for SARS-CoV-2 infection. Briefly, gramicidin S and melittin, were pre-incubated with SARS-CoV-2 virus for 1 hour at 37°C with 1.5 μg/100 μl and 3 μg/100 μl respectively. Later, the peptides containing the viral inoculum were used to infect Vero cells on the glass coverslips. After 3 hours of infection, the viral inoculum containing the peptides was replaced with fresh media with 10% FBS until 12 and 24 hrs. Parallel controls were maintained without the drug treatment. After 12 and 24 hours the treated and untreated cells were fixed with 4% paraformaldehyde and processed further for immunocytochemistry. The fixed samples were washed thrice with PBS and the cells were permeabilized using 0.3% Triton X-100 (Sigma, cat. no.: X100; Lot no.: 056K0045) in PBS for 15 min at room temperature (RT). Then the cells were washed with PBS, thrice for 5 min. each at RT. The cells were incubated with the 3% Bovine Serum Albumin (BSA) (Sigma) in PBS, for 1 hour at RT to block the nonspecific antibody binding. Later the experimental groups were incubated with the anti-sera for RBD of SARS-CoV-2 (1:200) prepared in 1% BSA made in PBS (anti-sera against SARS-CoV-2 was raised in rabbits and validated using ELISA at CCMB) overnight at 4°C. After incubation the cells were washed with PBST, thrice at RT for 10 min each. Later the cells were incubated with the secondary anti-Rabbit IgG antibody conjugated with Alexa Fluor 488 (Life Technologies, Cat. no.: A11008; Lot no.: 1735088) at the dilution of 1:200 in 1% BSA made in PBS. Rhodamine Phalloidin (Life Technologies, Cat. no.: R415; Lot no.: 1738179) was used to label F-actin. The cells were incubated with secondary antibody and Rhodamine phalloidin mix for 1 hour at room temperature. After incubation the cells were washed with PBST, thrice at RT for 10 min each. Then, the cover slips containing the cells were mounted over the pre-cleaned slides using Vectashield mounting medium containing DAPI (for nuclear staining) (Vector Laboratories, Cat. no.: H-1200, Lot no.: ZC1216). Images were obtained using confocal microscope FV3000 with software version 2.4.1.198 (Olympus Life Sciences Solutions) in Light Scanning Microscopy (LSM) mode.

### Proteomic Analysis

Total protein was extracted from the control Vero cells, Vero cells infected with SARS-CoV-2 and Vero cells infected with SARS-CoV-2 and treated with gramicidin S and melittin separately. The cells were collected at 24 and 48 hours independently. The samples were centrifuged and pellet was dissolved in protein solubilisation buffer [31, 32] and sonicated for 10 minutes at BSL3 lab facility. The protein samples were further centrifuged for 30 mins at 14000 RPM to remove the cell debris. Quantification of the pooled protein samples were performed using Amido Black method against BSA standard. A 200 μg of total protein from all the experimental groups were electrophoresed in 10% SDS-PAGE, Commassie R250 stained, destained and gel excised in to four fractions based on molecular weight. In-gel trypsin digestion and iTRAQ labeling and purification were performed as describe earlier [32–34]. iTRAQ label was labelled as 114 – Control; 115-Infection; 116-Infected cells treated with Melittin and 117 – Infected treated with Gramicidin S. All labelled peptides were pooled and purified by running through C18 column. Peptides were reconstituted in 5% acetonitrile (ACN) and 0.2% formic acid then subjected to the Liquid Chromatography Mass Spectrometry (LCMS/MSMS) analysis in OrbitrapVelos Nano analyzer (Q-Exactive HF). The proteomic data obtained from the mass spectrometer were analysed against human proteome and SARS-CoV-2 proteome data. All the obtained proteome data were tabulated and differential expression in SARS-CoV-2 proteins were estimated against the control negative samples. The obtained proteome data was analysed for its heat map expression profile using heatmapper software (www.heatmapper.ca). Network and pathway analysis of the associated proteins were performed using STRING v11.5.

### Molecular Docking

The receptor binding domain (RBD) of the SARS-CoV-2 spike protein was obtained by editing the crystal structure of the C-terminal domain of the SARS-CoV-2 spike protein in complex with human ACE2 (PDB id: 6zlg) [35]. The id of the structure used for gramicidin S monomer is CCDC 626343. Monomeric melittin structure was obtained by editing the crystal structure of tetrameric melittin (PDB id: 2mlt). The structures were generated using Discovery Studio 2019.

### Statistical Analysis

All the experiments were performed in duplicates with technical replicates (n=6). The data analysis and graphs were generated using GraphPad Prism (*Ver* 8.4.2). All the values were represented as mean ± SD.

## Acknowledgements

BKK would like to acknowledge financial support from Council of Scientific and Industrial Research (CSIR MLP0056). RN is Indian national Academy (INSA) Senior Scientist.

## Ethics approval statement

The Anti-SARS CoV-2 study was approved from Institutional Bio-safety Committee of CSIR-Centre for Cellular and Molecular Biology, Hyderabad, India.

